# Multiple modes of PRC2 inhibition elicit global chromatin alterations in H3K27M pediatric glioma

**DOI:** 10.1101/432781

**Authors:** James M. Stafford, Chul-Hwan Lee, Philipp Voigt, Nicolas Descostes, Ricardo Saldaña-Meyer, Jia-Ray Yu, Gary Leroy, Ozgur Oksuz, Jessica R. Chapman, Fernando Suarez, Aram S. Modrek, N. Sumru Bayin, Dimitris G. Placantonakis, Matthias A. Karajannis, Matija Snuder, Beatrix Ueberheide, Danny Reinberg

**Affiliations:** Department of Biochemistry and Molecular Pharmacology, NYUSoM, New York, NY, USA; Howard Hughes Medical Institute, Chevy Chase, MD, USA; Proteomics Laboratory, NYUSoM, New York, NY, USA; Laura and Isaac Perlmutter Cancer Center, NYUSoM, New York, NY, USA; Department of Neurosurgery, NYUSoM, New York, NY, USA; Kimmel Center for Stem Cell biology, NYUSoM, New York, NY, USA; Neuroscience Institute, NYUSoM, New York, NY, USA; Department of Pediatrics, NYUSoM, New York, NY, USA; Department of Patholog, Division of Neuropathology, NYUSoM, New York, NY, USA

**Author notes:** Wellcome Centre for Cell Biology, School of Biological Sciences, University of Edinburgh, Edinburgh, UK. Department of Pathology, Memorial Sloan Kettering Cancer Center, New York, NY USA. Department of Pediatrics, Memorial Sloan Kettering Cancer Center, New York, NY, USA. equal contributions (James M. Stafford, Chul-Hwan Lee).

## Abstract

A methionine substitution at lysine 27 on histone H3 variants (H3K27M) characterizes ~80% of diffuse intrinsic pontine gliomas (DIPG) and inhibits PRC2 in a dominant negative fashion. Yet, the mechanisms for this inhibition and abnormal epigenomic landscape have not been resolved. Using quantitative proteomics, we discovered that robust PRC2 inhibition requires levels of H3K27M greatly exceeding those of PRC2, seen in DIPG. While PRC2 inhibition requires interaction with H3K27M, we found this interaction on chromatin is transient with PRC2 largely being released from H3K27M. Unexpectedly, inhibition persisted even after PRC2 dissociated from H3K27M-chromatin suggesting a lasting impact on PRC2. Furthermore, allosterically activated PRC2 is particularly sensitive to K27M leading to a failure to spread H3K27me3 at distinct foci. In turn, levels of Polycomb antagonists such as H3K36me2 are elevated suggesting a more global, downstream effect on the epigenome. Together, these findings reveal the conditions required for H3K27M-mediated PRC2 inhibition and reconcile seemingly paradoxical effects of H3K27M on PRC2 recruitment and activity.

## INTRODUCTION

Histones form the core DNA packaging material in the nucleus. Even slight alterations in their amino acid composition can have dramatic consequences on chromatin structure affecting gene expression and genome integrity (*1*). One of the most striking examples is the lysine-to-methionine substitution at residue 27 on canonical histone H3.1 or its variant H3.3 (H3K27M). These mutations were discovered in ~80% of diffuse intrinsic pontine gliomas (DIPG), one of the most deadly and difficult-to-treat pediatric cancers (*2*). Despite affecting only one of 15 copies of the H3 genes, the presence of H3K27M significantly impairs the activity of PRC2, leading to a drastic reduction and re-distribution of di- and tri-methylation of H3K27 [H3K27me2 and -3; (*3–6*)] in the affected cells. These effects of H3K27M on PRC2, as well as those on the chromatin landscape have been linked to oncogenic transcriptional programs that give rise to the cancer stem cell-like and proliferative properties of DIPG (*4, 7, 8*). Therefore, understanding the underlying mechanisms of how H3K27M affects PRC2 and the global epigenetic landscape is imperative for understanding DIPG and ultimately, devising strategic ways to overcome the oncogenic potential of H3K27M.

Following the discovery of H3K27M, a hypothesis emerged suggesting that H3K27M traps and sequesters PRC2, thereby diminishing its activity (*3*). Supporting evidence came from the observation that PRC2 has a higher affinity for H3K27M relative to H3K27WT peptides and nucleosomes (*4, 9*). Moreover, structural studies revealed that H3K27M occupies the catalytic SET domain of EZH2, preventing the hydrolysis of the methyl donor S-adenosylmethionine (SAM) and the release of EZH2 (*3, 4, 9*). However, *in vivo* studies indicate that the genomic localization of H3K27M is often inversely correlated with PRC2 occupancy, challenging the idea that H3K27M sequesters PRC2 on chromatin (*6, 10, 11*). Notably, despite a drastic loss in H3K27me3 in affected cells, some loci retained PRC2 and H3K27me3 at abnormally high levels in H3K27M-expressing cells (*4, 5, 12*). Additional layers of complexity came from studies showing that H3K27M does not have consistent effects in all cell types (*7, 8*) nor is it able to strongly interact with PRC2 in all chromatin contexts (*6, 13*). Therefore, a model whereby H3K27M stably sequesters PRC2 on chromatin cannot explain all observations, indicating that further study is required to resolve these disparate observations and to uncover the mechanism(s) by which H3K27M affects PRC2 and the chromatin landscape in DIPG. This study aims to reconcile these findings by combining isogenic and patient-derived model systems to better understand the dynamic and nuanced impact of H3K27M on PRC2 while placing those effects in the molecular and broader chromatin context.

## RESULTS

### H3K27M to PRC2 ratio dictates the degree of PRC2 inhibition

Perhaps the most studied hypothesis for dominant negative inhibition of PRC2 by H3K27M is that H3K27M sequesters PRC2, preventing it from methylating the majority of H3K27WT nucleosomes (*3, 4, 9, 14*). This hypothesis relies on two assumptions: that H3K27M is more abundant than PRC2 and that PRC2 has a higher affinity for H3K27M-than H3K27WT-containing nucleosomes. Therefore, precise quantification of PRC2 relative to H3K27 methylation and H3K27M levels in DIPGs and comparable cell types is essential for understanding how the cellular stoichiometry of these factors contributes to cellular sensitivity to H3K27M. We hypothesized that the levels of PRC2 might render DIPG particularly sensitive to H3K27M given their profound loss in H3K27me2-3 relative to other brain tumors and cell types in the neural lineage (**Fig. 1A**).

**Figure 1.**
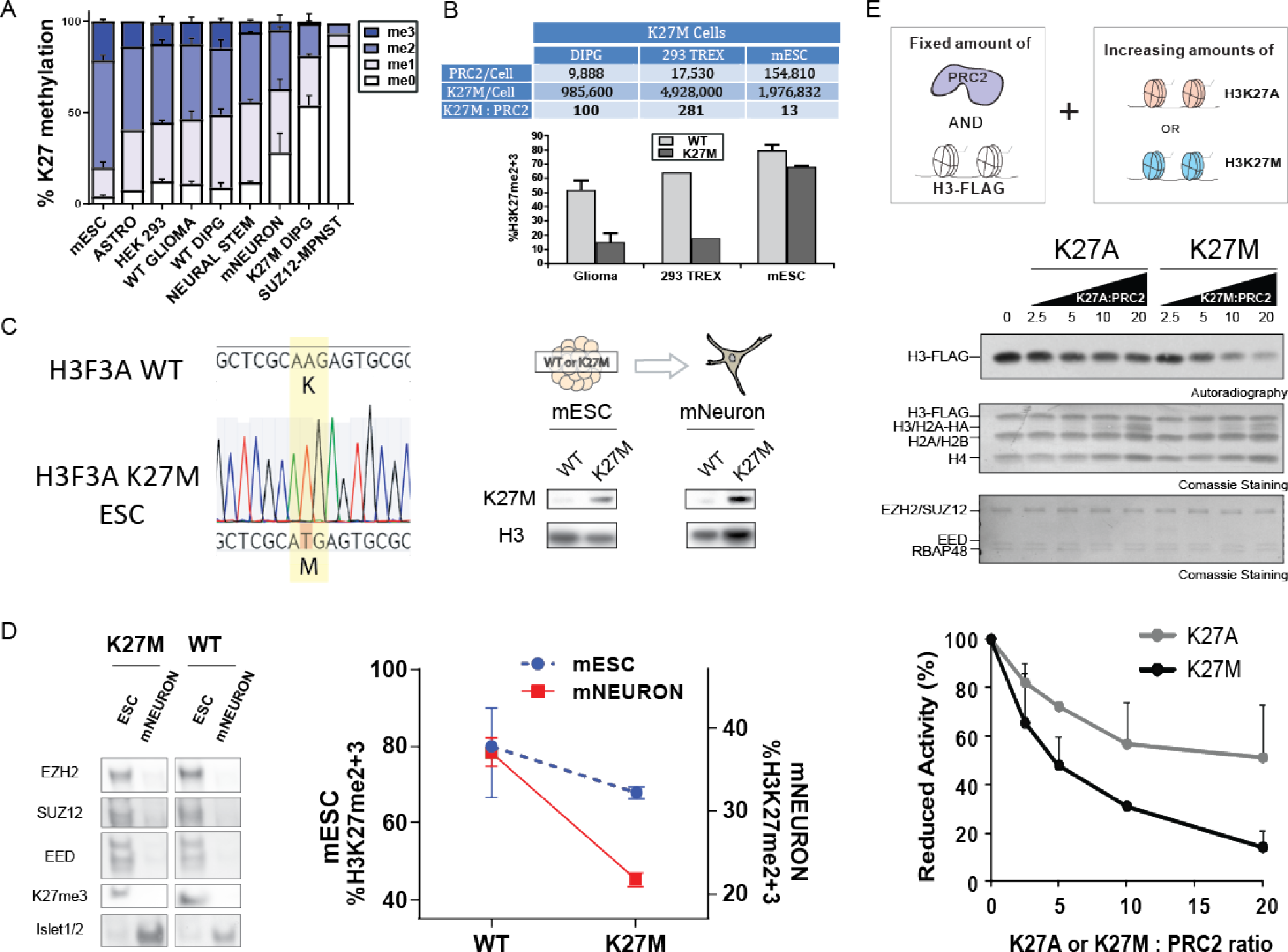
PRC2 inhibition as a function of the H3K27M to PRC2 ratio. **(A)** Samples are ranked left-to-right based on levels of H3K27me2/−3 as detected by mass-spectrometry. [mESC: mouse embryonic stem cell (*n=2)*, ASTRO: human astrocyte (*n=1)*, HEK 293 (*n=2*), WT GLIOMA: H3K27WT cortical glioma (*n=4)*, WT DIPG: H3K27WT DIPG (*n=2)*, NEURAL STEM: human neural stem cells (*n=2)*, mNEURON: mouse motor neuron *(n=3)*, K27M DIPG: DIPG with H3K27M (*n=2* for H3.1K27M and *n=2* for H3.3K27M), SUZ12 MPNST: Malignant peripheral nerve sheath tumor that is SUZ12 null (*n=*1)]. **(B)** PRC2 molecules per cell (average of EED, EZH2, and SUZ12) were determined by quantitative mass spectrometry (MS) and are presented in the table along with the relative ratio of H3K27M to PRC2. Levels of H3K27me2/−3 determined by MS are presented in the chart below. DIPG and 293 TREX cells with a large excess of K27M to PRC2 showed the most robust attenuation of H3K27me2/−3 levels while embryonic stem cells (mESC) with a more modest excess of K27M (~13 fold) showed less robust loss in K27me2/−3 relative to their WT counterparts (see **Table S1** for cell lines details and **fig. S1** for cell lines used in PRC2 quantitation analyses). **(C)** mESC generated using CRISPR harboring either WT or a K27M mutation at H3F3A were differentiated to motor neurons (mNEURON). *Left*, Sanger sequencing results for H3F3A K27M mESCs compared with WT mESCs. *Right*, Western blot validating the cell lines by H3K27M protein expression. **(D)***Left*, Western blot validating the cell lines and increased K27M expression with significantly reduced levels of PRC2 core components in mNEURON. Islet1/2 served as a motor neuron marker. *Right Graph*, Differentiation to mNEURON led to decreased K27me2/−3 in K27M cells, relative to their mESC precursors, as measured through MS (*n=2*/cell type). **(E)***Top*, Histone methyltransferase assays (HMT) containing PRC2 and increasing ratios of 8x-oligonucleosomes comprising H3K27A or H3K27M and HA-tagged H2A. Substrate oligonucleosomes were distinguishable by their reconstitution with H3-FLAG. *Middle*, Representative HMT assay shows levels of methylation and relative concentration of each HMT component. *Bottom*, Graphs quantitate the relative amount of ^3^H-SAM incorporate into histone H3-FLAG. Higher H3K27M:PRC2 ratios produce larger deficits in PRC2 activity (*n=3*/data point). Data Plotted as Mean±SD.

Testing this hypothesis required a quantitative measure of PRC2 molecules per cell, leading us to develop a mass spectrometry (MS) parallel reaction-monitoring assay (*15*). Briefly, PRC2 core proteins (EZH2, EED, and SUZ12) were quantified with a targeted method using heavy isotope-labeled synthetic peptides as internal standards (**fig. S1A-S1C**). Our approach revealed a wide range of PRC2 levels across different cell types, with CRISPR-generated H3K27M (on H3F3A) E14 mouse ESCs (mESC) containing relatively high PRC2 levels (~150,000 PRC2 molecules/cell) (**Fig. 1B** **and** **fig. S1D**). In our isogenic mESC and 293 TREX system below we chose H3.3K27M as it is the most frequently occurring H3K27M mutation (*2*). Interestingly, H3K27M DIPG (both H3.1- and H3.3K27M DIPGs) were at the lower end of the spectrum (~5-10,000 PRC2 molecules/cell; **fig. S1D**). By comparing the levels of PRC2 with those of H3K27M as determined by relative quantitative histone MS, we observed that H3K27M-bearing DIPG exhibited ~100-fold excess of H3K27M over PRC2 (**Fig. 1B** and see **Table S1** for details on cell lines). In accordance, H3K27M-bearing DIPGs showed a robust loss in H3K27me2-3 deposition relative to WT DIPG (**Fig. 1B**). Similarly, 293 TREX cells with inducible expression of H3K27M (H3.3K27M) also showed a large excess of H3K27M relative to PRC2 (over 200-fold) and a substantial deficit in H3K27me2-3. However, mESC harboring H3.3K27M showed only a modest loss in H3K27me2/−3, relative to WT mESC. Given that H3K27M was only in ~10-fold excess of PRC2 in mESC, the results suggested that H3K27M must be largely in excess of PRC2 to achieve the robust decrease in H3K27me2-3 seen in DIPG and other cell types. Importantly, these results are consistent with previous speculation that high amounts of a K-to-M mutation is needed to inhibit various histone methyltransferases (*3, 16*).

To test whether shifting the relative abundance of H3K27M and PRC2 during lineage commitment could alter sensitivity to H3K27M inhibition, we differentiated H3.3K27M mESC as described above into cervical motor neuron precursors [mNEURON; (*17*)] (**Fig. 1C**). Our quantitative mass-spec revealed that PRC2 decreased as cells proceeded down the motor neuron lineage to levels consistent with those seen in DIPG [5-10,000 molecules/cell; **fig S1**]. Importantly, we observed that K27M produced a greater loss of H3K27me2/−3 in mNEURON than was seen in undifferentiated mESC (**Fig. 1D)**. Although we cannot rule out the contribution of an unknown factor(s) during the differentiation process, these results together with those analyzing the quantitative H3K27M:PRC2 ratio in different cells, demonstrate that the critical force for the H3K27me3 reduction is due the large excess of K27M over PRC2. These observations are consistent with K27me2/−3 loss when the H3K27M:PRC2 ratio is high, which we tested directly using a histone methyltransferase assay (HMT) in a fully biochemically reconstituted system (**Fig. 1E**). The addition of increasing amounts of recombinant oligonucleosomes comprising H3K27M gave rise to heightened sensitivity of PRC2 to inhibition, while the control using oligonucleosomes where lysine-27 was mutated to alanine (H3K27A) were ineffectual (**Fig. 1E**). Together, the quantitative approach measuring H3K27M:PRC2 ratio in different cell lines including patient-derived DIPG model systems, the *in vitro* differentiation studies together with the biochemical approach, revealed that cells containing low amounts of PRC2, relative to H3K27M such as DIPG, motor neurons and 293 TREX are sensitive to H3K27M-mediated inhibition of PRC2 activity.

### PRC2 is transiently recruited to H3K27M-containing chromatin

Given the high excess of H3K27M to PRC2 in DIPG (**Fig. 1B**) and the higher affinity of PRC2 for H3K27M *in vitro*, H3K27M would be expected to sequester all PRC2 in a DIPG cell (*3, 9, 16*). However, the following observations argue against this assumption: 1) PRC2 and H3K27M often do not co-localize on chromatin under some conditions of steady-state H3K27M expression, as measured by ChIP (*6, 10, 18*) and 2) H3K27me3 can be retained at specific loci as well as redistributed in H3K27M cells (*4–6, 10, 12*). Reasoning that the steady-state dynamics between H3K27M and PRC2 reflect only the outcome of this dysregulated process, we sought a means to detect changes in the interaction between H3K27me3 and PRC2 and subsequent effects on PRC2 as a function of the rate of H3K27M deposition. To this end, we used an inducible expression system in 293 TREX cells to monitor the early and later events following H3K27M expression (**Fig. 2A)**. This inducible system recapitulated the global loss of H3K27me2/−3 seen in H3K27M-bearing DIPG (**Fig. 2B & 1A**). Spike-in normalized ChIP-seq experiments across this time course revealed that PRC2 primarily co-localized with H3K27M at early time points, namely after 6 and 12 hr of expression, but was largely excluded from H3K27M sites at the late, 24 hr time point (**Fig. 2C left panel**). Genome-wide analysis indicated that there are nearly 1,500 PRC2 occupied domains unique to the 6 and 12 hr time points that are not present at 24 hr (**Fig. 2C right panel)**. Importantly, those early PRC2 loci are only seen in H3K27M, but not H3K27WT-expressing cells and co-localized with H3K27M (**Fig. 2C)**. Combined with the higher affinity of PRC2 for H3K27M nucleosomes [(*4*) and see below] and peptides (*9*), our results demonstrated that H3K27M-containing nucleosomes transiently recruit PRC2 to loci not typically occupied by PRC2 in a wild-type setting, leading to aberrant deposition of PRC2 early after H3K27M expression (**Fig. 2**).

**Figure 2.**
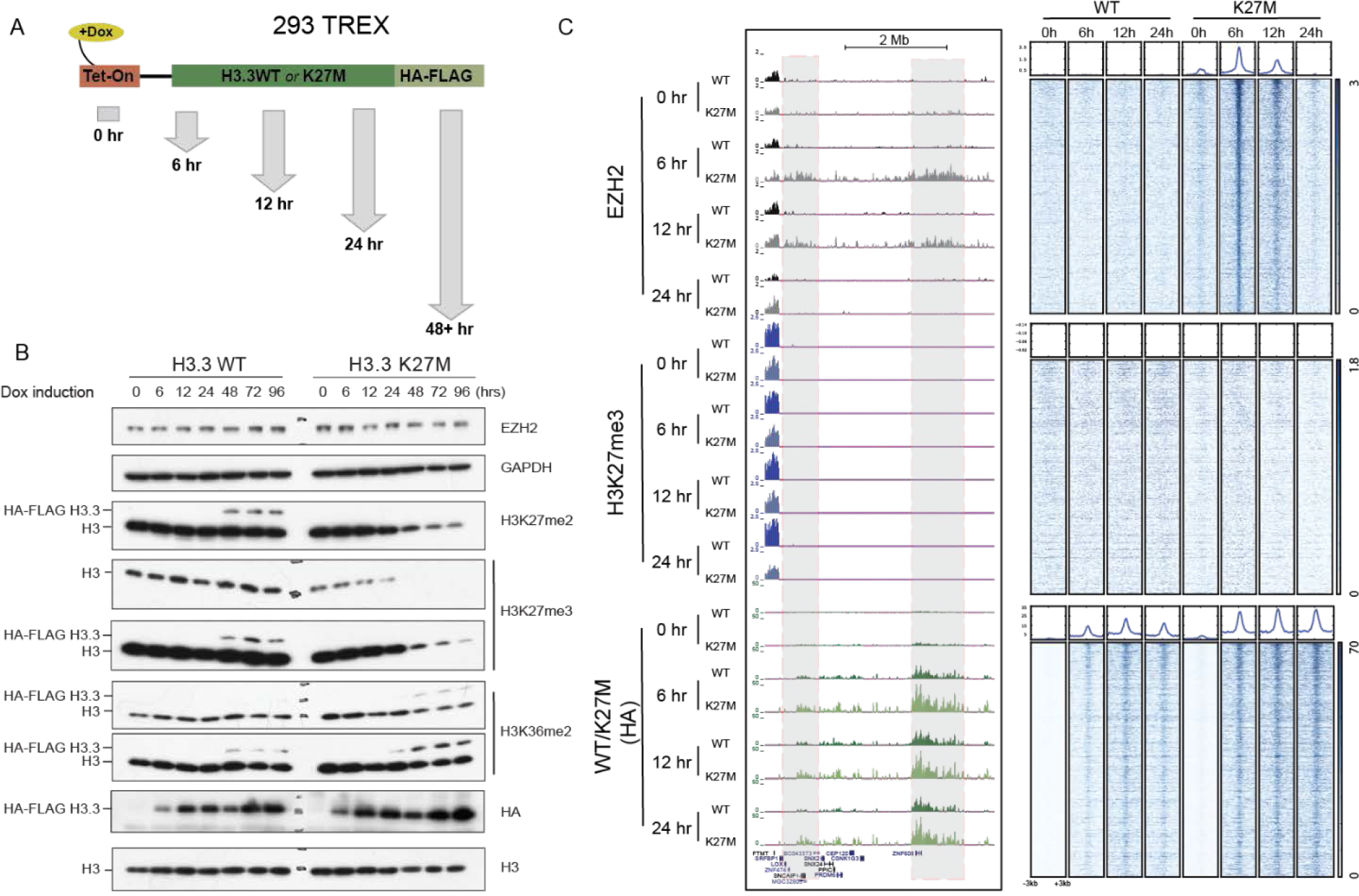
Abnormal recruitment and release of PRC2 from H3K27M chromatin. **(A)** 293 TREX cells were induced to express C-terminal, FLAG-HA tagged-H3.3K27M (K27M) or -H3.3K27WT (WT) during the indicated time course. **(B)** Western blots revealed the progressive loss in H3K27me2/−3 and gain of H3K36me2 as a function of time of H3.3K27M expression (note slower migration of tagged H3.3 versions). **(C)** ChIP-seq for EZH2, H3K27me3 and HA-tagged WT/K27M histones performed at select time points and displayed as representative enrichment tracks (*left*). EZH2 peaks detected exclusively under the 6 and 12 hr conditions in K27M-expressing cells are highlighted in gray. *Right* – Heatmaps were generated by centering and rank-ordering the 6 hr EZH2 peaks detected within EZH2 regions present only at the early 6 and 12 hr time points (1,450 domains; 2,325 peaks). Corresponding average density profiles are plotted at the top of the heatmaps. Peaks occurring specifically at the early time points (6 and 12 hr time points) strongly co-localize with K27M, are unique to the K27M cells and are depleted by 24 hr.

Given the transient nature of PRC2 recruitment to H3K27M, numerous factors might contribute to its eviction including the known PRC2 antagonist, H3K36 methylation, especially when it occurs directly adjacent to H3K27M (*i.e., in cis*). Indeed, western blot analysis of extracts derived from the inducible 293 TREX system revealed enrichment of H3K36me2 on H3.3K27M histones, supporting their co-occurrence *in cis*, which coincided with a loss of H3K27me2/−3 (**Fig. 2B**). This was particularly evident in H3K27M-bearing DIPG where mass-spectrometry revealed that H3K36me2 was overwhelmingly the predominant modification adjacent to H3K27M regardless of whether it was H3.1- or H3.3K27M (**fig. S2A)**. Importantly, in our time course studies, a strong co-occurrence *in cis* of H3K36me2 and H3K27M emerged at 24 hr at which time EZH2 occupancy at H3K27M-containing loci is lost, relative to earlier time points (**Fig. 2B**). Therefore, eventual PRC2 eviction and stable exclusion from K27M loci might be related to H3K36me2 deposition, presumably by preventing PRC2 interaction with the H3 N-terminal tail (*19–22*). Other antagonistic modifications such as histone acetylation might also contribute as has been suggested by others [(*6, 13*) and *see our data below*]. In a broad sense, the deposition of H3K27M largely into transcriptionally active regions in both the isogenic K27M 293 TREX and K27M DIPG cells themselves (**fig. S2B**), suggests a dynamic environment wherein histone modifications associated with transcription activity [*this study* and (*6, 13*)], histone turnover (*23*) and even nascent mRNA might lead to the failure of K27M to stably recruit PRC2 [i.e., the eviction of PRC2 (*11, 24*)]. While we cannot exclude the possibility that the release of PRC2 from H3K27M arises from a more direct effect on PRC2 itself (see Discussion), we expect that future studies will find that the interaction and release of PRC2 from H3K27M is determined by the interplay between the local chromatin environment and structural changes to PRC2 itself.

### Persistent inhibition of PRC2 after its dissociation from H3K27M

We next sought to better understand the impact of H3K27M on PRC2 activity at more typical PRC2 target sites. First, we identified 616 broad PRC2 (EZH2) domains present in both H3K27M and H3K27WT cells across all time points and found that despite the retention of EZH2 at a subset of genes at 24 hr in H3K27M cells, a global loss of H3K27me3 was evident (**Fig. 3A** **and** **fig. S3** for visualization of the broad domains). As the cells progressed attaining steady-state levels of H3K27M, the loss in H3K27me3 was even greater in magnitude, despite the retention of EZH2 and the lack of H3K27M enrichment at those same sites after 72 hr of H3K27M expression (**Fig. 3B**). Together, these results are consistent with recent work showing that loci depleted of H3K27me3 in H3K27M cells have equal PRC2 occupancy, as well as emerging evidence that PRC2 recruitment can be uncoupled from its enzymatic activity [(*5, 25*) and see below].

**Figure 3.**
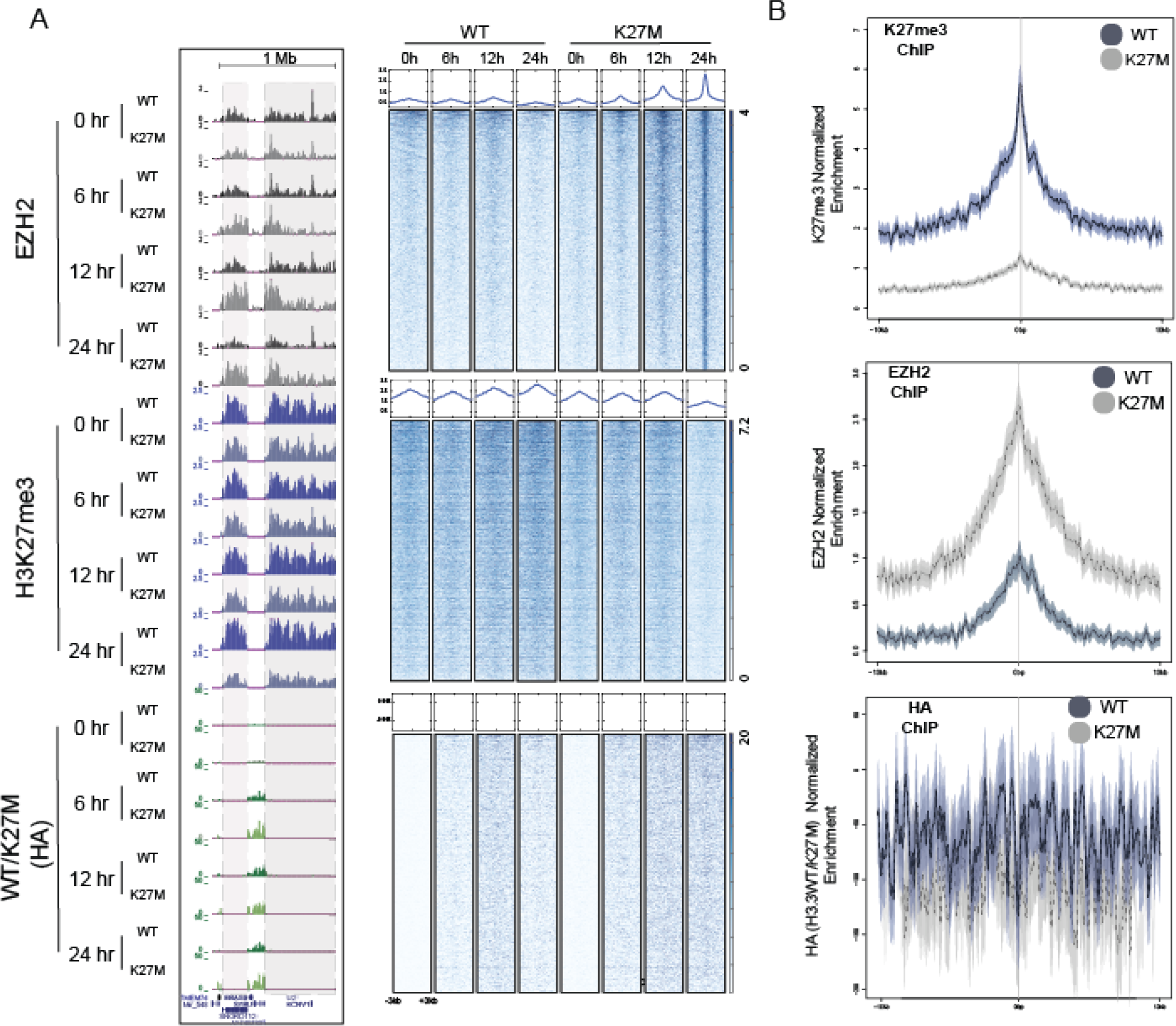
PRC2 is recruited to normal target sites but is progressively less active on chromatin following K27M expression. **(A)** ChIP-seq for EZH2, H3K27me3 and HA-tagged WT/K27M histones performed at select time points and displayed as representative enrichment tracks (*left*). PRC2 (EZH2) domains detected in both WT- and K27M-expressing cells across all time-points are highlighted in gray. *Right*-Heatmaps centering on the 24 hr EZH2 peaks within the common PRC2 domains (616 domains; 2,253 peaks) indicate strong EZH2 enrichment at that time point, decreased H3K27me3 and little detectable co-enrichment with H3K27M (HA). **(B)** To confirm the effects at the 24 hr time point, ChIP-seq was performed after 72 hr of H3.3K27M (K27M) and H3.3K27WT (WT) expression for EZH2, HA, and H3K27me3. Genome-wide enrichment plots centered on the common EZH2 domains corresponding to those in *Panel A* are presented. Despite a loss in H3K27me3 in H3K27M cells relative to WT (*top*), EZH2 occupancy is increased and the tagged histones are not enriched (*middle* and *bottom*, respectively).

We next investigated whether the interaction of PRC2 with H3K27M-containing nucleosomes might have a lasting impact on its function. This was motivated by our observations that; 1) newly expressed K27M deposits at sites that are not usually regulated by PRC2 and PRC2 is transiently recruited at these new sites, as well as, 2) some PRC2 regulated genes maintain PRC2, but its activity is reduced in H3K27M cells (**Fig. 2** **and** **3**). To study this, we expressed tagged-PRC2 in H3K27M- and H3K27WT-expressing 293 TREX cells (**fig. S4A** **and B**). PRC2 was then purified using a combination of conventional and tandem-affinity chromatography yielding PRC2 in correct stoichiometry between samples (**Fig. 4A**). Most importantly, H3K27M was not detected by western blotting and mass spectrometry analysis in the purified PRC2 fractions (**Fig. 4A** **and** **Fig. S4C**). Interestingly, we found that the catalytic activity of PRC2 purified from H3K27M-expressing cells was markedly decreased relative to PRC2 purified from H3K27WT cells and this inhibition was SAM dependent (**Fig. 4B**) which is consistent with the previous observation demonstrating that SAM is necessary for inhibition of the methyltransferase by K-to-M mutation on histone H3 (*26*). This lasting H3K27M inhibitory effect on PRC2 was also observed using a system reconstituted with all recombinant components, including PRC2 comprising only its 4 core subunits (EZH2, EED, SUZ12 and RBAP48; **Fig. 4C**) and oligonucleosomes formed with bacterially-expressed histones (**Fig. 4D**). After release from H3K27M-containing oligonucleosomes, the resultant PRC2 was generally less active (**Fig. 4E**), in line with a wide-spread attenuation of H3K27me2/−3 in K27M DIPG. Given that lasting PRC2 inhibition is seen *in vivo* as well as within a fully recombinant system, we speculate that H3K27M-containing chromatin alters the core PRC2 complex through a stable conformational change (see Discussion).

**Figure 4.**
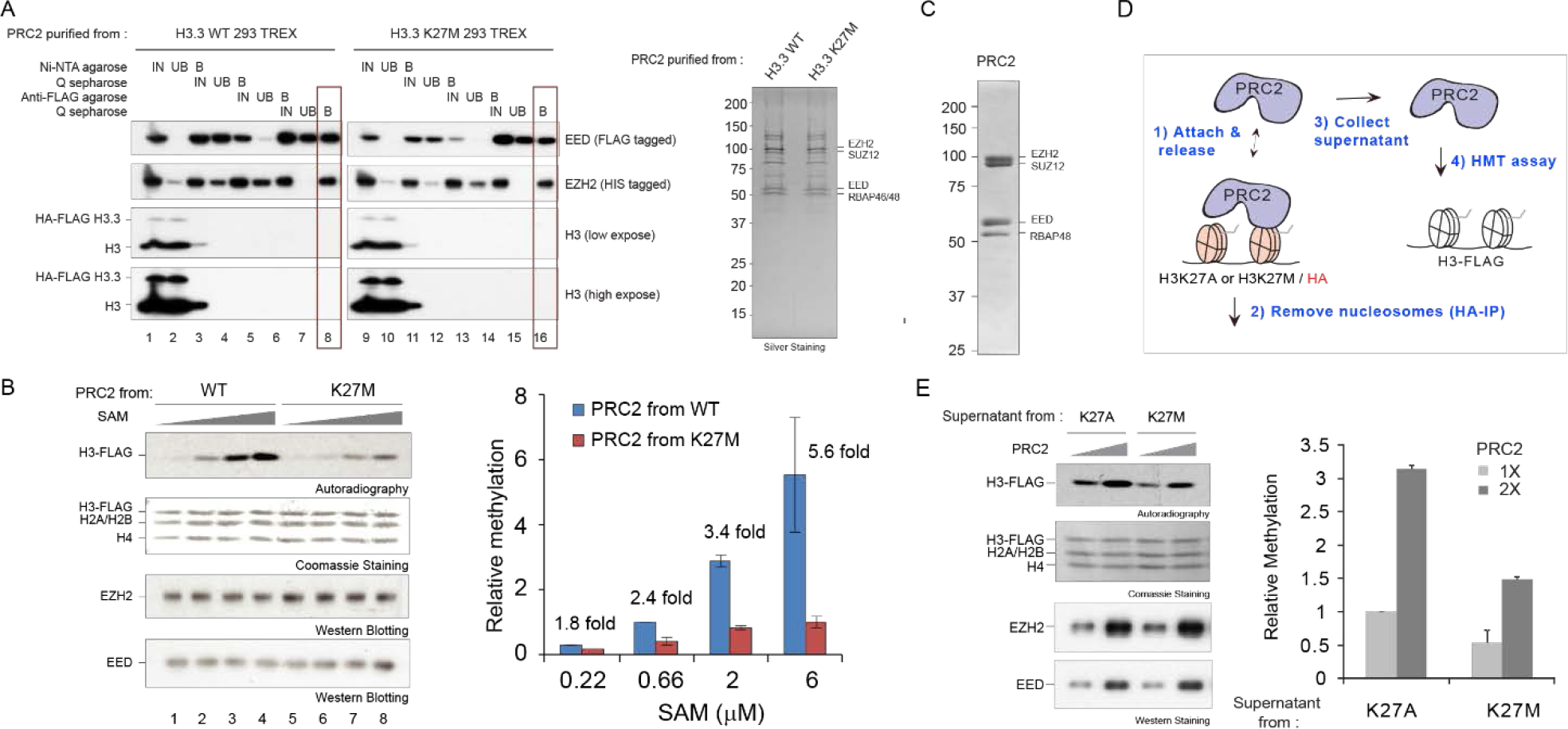
PRC2 purified from K27M cells or nucleosomes is less active. **(A)** PRC2 containing 6xHIS-tagged EZH2 and FLAG-tagged EED was purified from 293 TREX cells expressing H3.3K27M or H3.3K27WT for 3 days. *Left*, Western blot of levels of EZH2, EED, and histone H3 at each purification step. Note that histones and non-stoichiometric levels of EED or EZH2 were well-removed through purification (IN = Input, UB = Unbound, B = Bound; see also Fig. S6). Lanes 8 and 16 (boxed) represent the final, purified PRC2 complex used in subsequent assays. *Right*, Silver staining showing the relative purity of each purified PRC2 complex. **(B)** *Left*, Purified PRC2 complexes as in *Panel A* were subjected to HMT assays using 8x-oligonucleosomes with H3-FLAG as substrate (300 nM) and increasing amounts (0.22, 0.66, 2, and 6 μM) of SAM (^3^H-SAM:SAM=1:9). *Right*, Quantification of the relative amounts of ^3^H-SAM incorporated into histone H3-FLAG substrate (*n=2*). Data Plotted as Mean±SD. **(C)** Coomassie blue staining shows recombinant PRC2 (EZH2, EED, SUZ12 and RBAP48) purified from SF9 cells. **(D)** Schematic representation of the method used to recover recombinant PRC2 after its association with different types of recombinant chromatin. Recombinant PRC2 (15 and 30 nM) was initially incubated with 8x-oligonucleosomes containing HA-tagged H2A and either H3K27A or H3K27M (300 nM) for 1 hr at 30 °C. PRC2 was then recovered by HA immunoprecipitation and the supernatant was collected. Equal amounts of unbound PRC2 (1x or 2x) was incubated with ^3^H-SAM (500 nM) using 8x-oligonucleosomes comprising H3-FLAG as substrate (300 nM). **(E)** A representative image of the HMT assays (*Left*). Quantitation of relative amount of ^3^H-SAM incorporated into histone H3-FLAG substrate (*n=3; Right*). Data Plotted as Mean±SD.

### Inhibition of allosteric activation of PRC2 and spreading of H3K27me3

We noted a strong similarity between the phenotype of H3K27M cells in which PRC2 is recruited but exhibits diminished activity and that of PRC2 mutants that are deficient in allosteric activation (*25*). Thus, we next tested whether H3K27M may preferentially impair allosteric activation of PRC2. To accomplish this, we incubated PRC2 with increasing amounts of the allosterically activating H3K27me3 peptide in the presence of SAM and nucleosomes comprising either H3K27M or control H3K27A (HA-tag on H2A; **Fig 5A**). PRC2 activity was subsequently assessed by adding WT nucleosomes (distinguished by a C-terminal FLAG-tag on H3; **Fig. 5A**). Strikingly, while there was little difference in the absence of H3K27me3 peptide, PRC2 was refractory to stimulation by H3K27me3 in the presence of H3K27M nucleosomes (**Fig. 5B)**. Furthermore, the presence of H3K27me3 selectively increased the binding affinity of PRC2 towards H3K27M-containing chromatin (**Fig. 5C**). Combined, these results indicate that the allosterically activated form of PRC2 (*25, 27*) binds stronger to H3K27M and that PRC2 catalysis is more sensitive to H3K27M.

**Figure 5.**
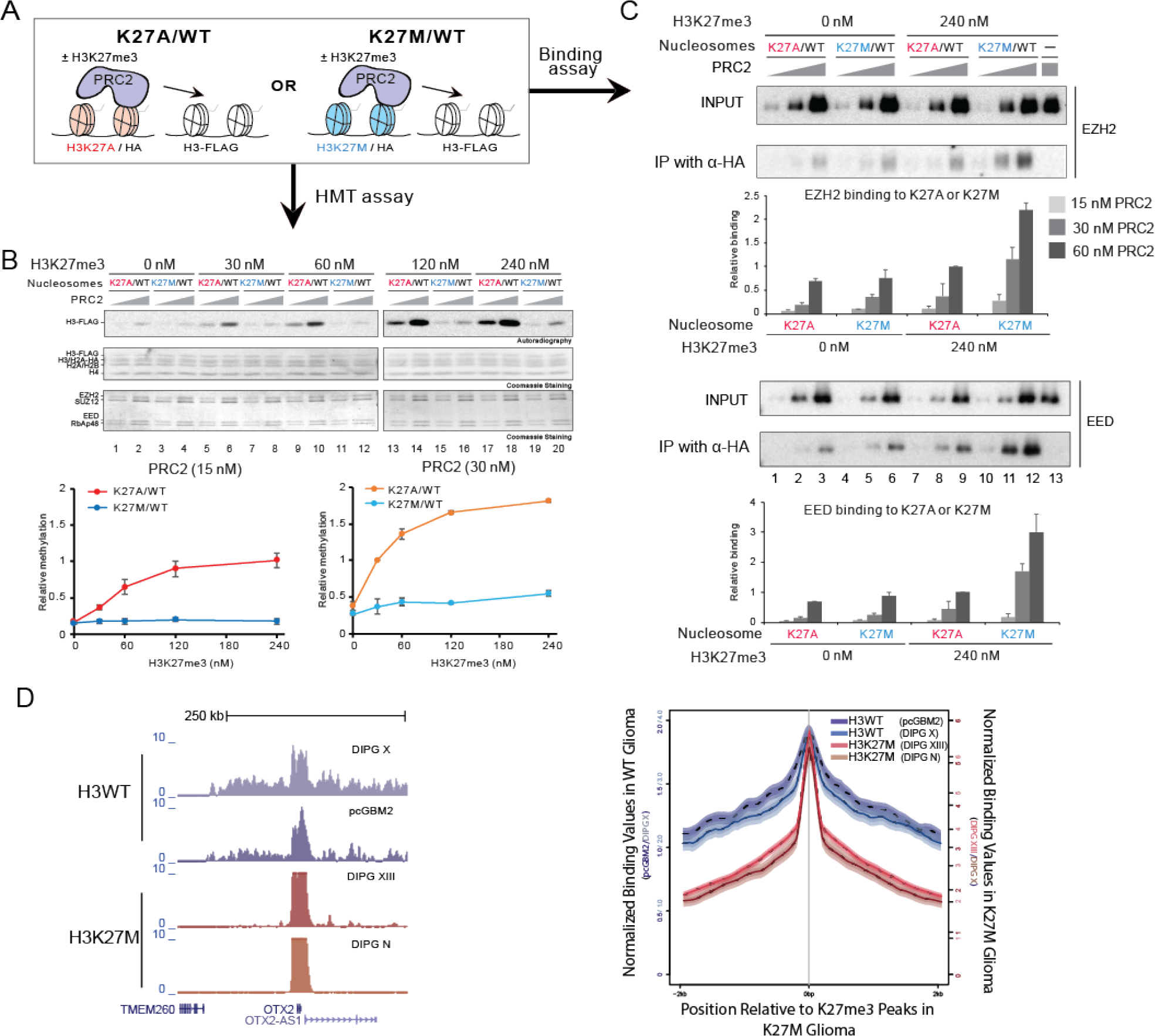
H3K27M preferentially inhibits allosterically activated PRC2 *in vitro* and restricts remaining H3K27me3 in DIPG. **(A)** Schematic representation of experiments in which PRC2 was incubated with ^3^H-SAM, increasing concentrations of H3K27me3 peptide and 12x-oligonucleosomes comprising HA-tagged H2A and either H3K27A or H3K27M. Oligonucleosomes containing FLAG-tagged-H3 (300 nM) were added as substrate. **(B)** One half of the reaction was subject to autoradiography with subsequent quantification of the relative amount of ^3^H-SAM incorporated into the FLAG-tagged histone H3 (: *n=3*/data point). Data Plotted as Mean±SD. **(C)** The other half of the reaction was used to measure the amount of PRC2 retained on H3K27M-versus H3K27A-containing 12x-nucleosomes by western blot for EZH2 and EED after immunoprecipitation of the HA-tagged-H2A-containing oligonucleosomes. The band intensity of EZH2 or EED served as a readout of EZH2 bound or EED bound on H3K27M- or H3K27A-containing 12x-oligonucleosomes, as indicated *(n=3/data point*). Data Plotted as Mean±SD. **(D)** Representative H3K27me3 ChIP-seq tracks from patient-derived glioma cell lines with K27M (*n=2* cell lines) or WT H3 (*n=2* cell lines) showing very sharp, narrow peaks in the K27M glioma and much broader peaks in the WT glioma (*Left*). Genome-wide trend plot indicating that the remaining K27me3 peaks in the K27M glioma are more punctate in contrast to the WT glioma which show large, broad H3K27me3 signal indicative of typical polycomb domains (*Right*).

To examine whether a similar defect in catalysis exists in H3K27M-bearing glioma, we used an internal spike-in normalized ChIP-seq protocol and directly compared H3K27me3 deposition in two independent H3K27M DIPG and H3K27WT glioma cell lines (selected based on being pediatric WHO Grade IV, matched culture conditions and IDH1 wild-type). Interestingly, the H3K27me3 peaks in the H3K27M cells appeared sharper and larger in magnitude than those observed in WT cells (**Fig. 5D**-**left**). Genome-wide analysis confirmed these observations, and more importantly revealed that the “remaining” H3K27me3 peaks in H3K27M glioma were not subject to spreading into large repressive H3K27-me-containing domains as observed in WT cells, a process likely dependent on PRC2-mediated allosteric activation (*25, 27*) (**Fig. 5D-right**). Our results are consistent with a broader mechanism of K-to-M substitutions as evidenced in *S. pombe* wherein H3K9M induces a gain in H3K9me3 and its methyltransferase (Clr4) at H3K9me3 nucleation sites, while H3K9me3 spreading across domains is lost (*16*). Our results account for the recent descriptions of H3K27M isogenic and glioma cells wherein there is a focal gain in H3K27me3 at previously established PRC2 target loci that are CpG island-dense (*5, 7, 8, 12*). Thus, by impairing allosteric activation of PRC2 and thus spreading of H3K27me3 (*25, 27*), H3K27M achieves both a global loss of H3K27me3 and yet, a focal accumulation of H3K27me3 at PRC2 recruitment sites.

### Altered histone modification landscape in H3K27M DIPG

Given the widely documented cross-talk among histone modifications, we tested whether the presence of H3K27M might affect the global histone modification landscape that could further affect PRC2 activity, resulting in an aberrant epigenetic profile in DIPG. In H3K27M-bearing DIPGs, a few histone modifications such as H3K27 acetylation have already been shown to increase in concert to the decrease in H3K27me3 (*3, 4, 12, 28*). However, a systematic analysis of the histone modification profile of DIPGs has so far been hampered, perhaps by the scarcity of suitable WT controls given that the majority of DIPGs carry the H3K27M mutation. Here, we were able to utilize the same quantitative histone modification MS approach as above on a panel of human primary H3K27WT glioma lines, two different H3K27WT DIPGs and four different H3K27M DIPGs to identify global histone modification alterations against a relevant control background. Consistent with other studies was a large-scale gain of histone acetylation in H3K27M DIPG (**fig. S5**). However, we observe that the largest gain of acetylation is on the H4 tail (~20% gain). This finding suggested that histone acetylation events beyond the relatively smaller net gain in H3K27 acetylation (~ 0.3%) reported here (**fig. S5**) and elsewhere (*3, 6, 29*) might be important drivers of the disease.

The most striking result was a significant gain in H3K36me2 within H3K27M DIPG cells (**Fig. 6A**). ChIP-seq indicated higher levels of H3K36me2 on H3K36me2-target genes in K27M DIPG relative to WT (**Fig. 6B and C**). In our isogenic H3K27M 293 TREX system, we further observed an invasion of H3K36me2 largely into former PRC2 domains that lost H3K27me3 (**Fig. S6**). These alterations in the euchromatic H3K36me2-domains and the corresponding heterochromatic H3K27me3-domains are reminiscent of sarcoma cells that express H3K36M and exhibit dampened H3K36 methyltransferase(s) activity leading to abnormal deposition of H3K27me3 (*30*). Due to the heterogeneity of pediatric glioma cell culture lines, we selected a WT (DIPG X) and a K27M (DIPG XIII) cell line to compare by RNA-seq as these are better matched with respect to both being DIPG, as well as their growth properties as non-adherent tumorspheres. As expected, the regions that gain H3K36me2 in K27M DIPG also show upregulation of those genes (**fig. S7A**) and are associated with proneuronal phenotypes, cancer pathways and cell adhesion phenotypes seen in H3K27M DIPG by KEGG pathway analysis (**fig. S7B**). Thus, in each of these cases, the K-to-M mutations disrupt heterochromatic and euchromatic domains that have potential to drive aberrant gene expression patterns important for proliferation, cancer stem cell properties, and oncogenesis (*4–7*).

**Figure 6.**
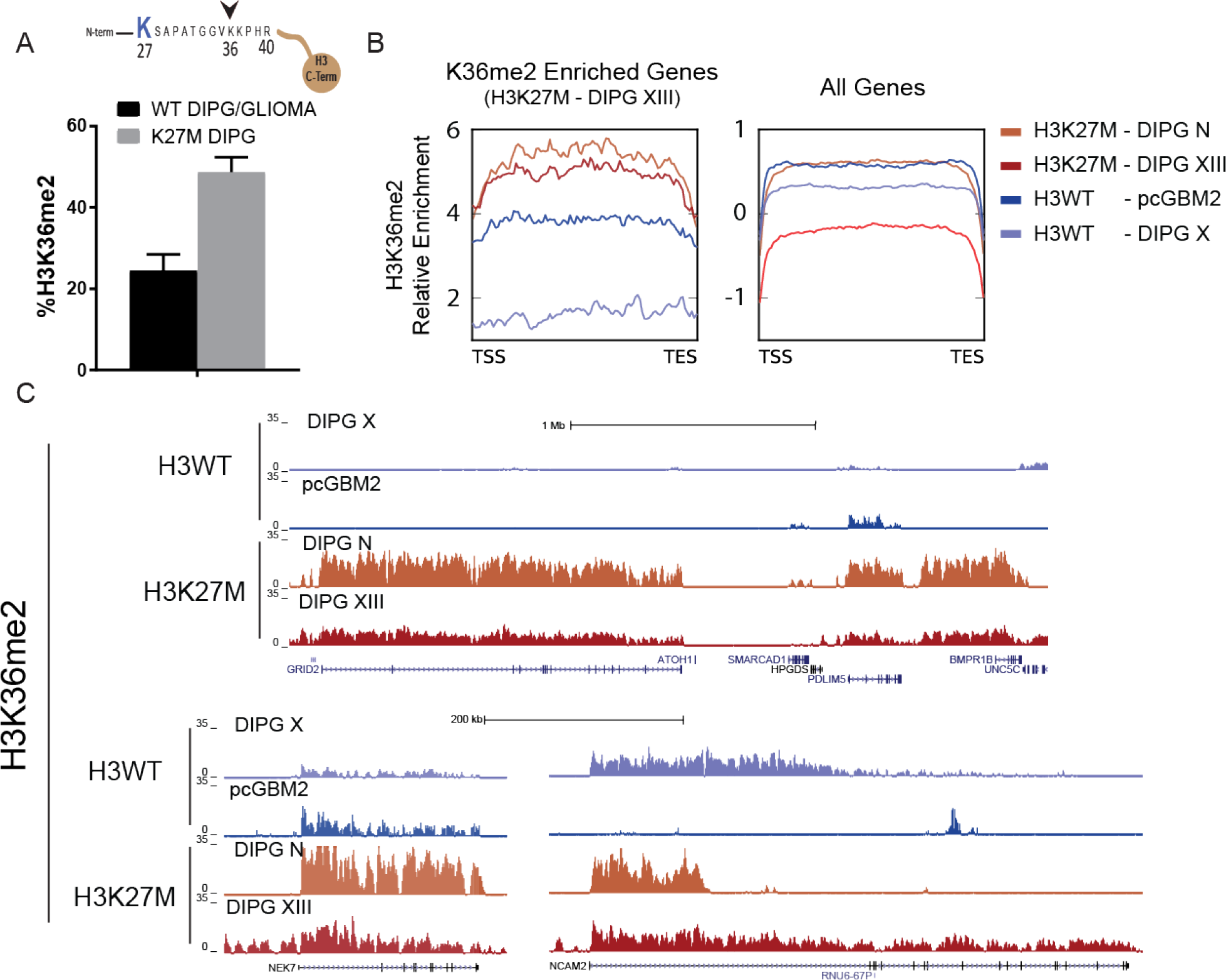
Gain of K36me2 in K27M DIPG. **(A)** H3K36me2 levels are significantly higher in H3K27M DIPG (*n=2* for H3.1K27M and *n=2* for H3.3K27M) than in WT DIPG/GLIOMA (H3K27WT cortical glioma *n=4*, H3K27WT DIPG *n=2)*, as revealed by MS (see *Fig. S7*). Data Plotted as Mean±SEM. **(B)** H3K36me2 ChIP-seq performed with glioma cell lines bearing either WT histone H3 (*n=2* total) or H3K27M, as indicated (*n=2*; see Table S1 for details). The *left* trend plot was generated by first identifying genes that displayed significant H3K36me2 enrichment above input for the H3K27M-bearing DIPG XIII cell line (1,727 genes). Subsequently, average H3K36me2 levels were plotted for each glioma cell line across those genes (transcriptional start to end site). As a control, H3K36me2 was plotted over all genes in the genome (*right*). Combined, H3K36me2 enriched genes in the H3K27M-DIPG showed overall higher levels of H3K36me2 relative to glioma with WT histone H3. **(C)** Representative ChIP-seq tracks show generally elevated levels of H3K36me2 across these genes in H3K27M DIPG. Variations of the effect can be seen in these gliomas: some genes devoid of H3K36me2 in H3WT gain H3K36me2 in H3K27M, while others contain H3K36me2 in both H3WT and H3K27M glioma, yet the levels in the H3K27M lines are generally higher.

## DISCUSSION

A primary challenge in the H3K27M field has been to resolve the contradictory observations indicating that PRC2 has a high affinity for H3K27M peptides, yet PRC2 and H3K27M are often mutually excluded from chromatin in K27M DIPG (*4, 6, 9, 10*). Our studies revealed that interaction of H3K27M with PRC2 is a dynamic process that cannot be captured by static, steady-state approaches. Namely, there is an initial phase after H3K27M is expressed and incorporated into chromatin, followed by PRC2 recruitment to H3K27M-containing chromatin, presumably due to its higher affinity towards H3K27M (Fig. 2C). However, in the next phase, PRC2 is released from H3K27M as they do not co-localize at steady state conditions in both isogenic 293 TREX systems (e.g., this study, Fig. 2 & 3) as well as in the H3K27M DIPG themselves (*6*). This dynamic model therefore accommodates both the finding of high H3K27M and PRC2 affinity in select assays as well as their failure to stably colocalize on chromatin in cells. In line with the more fluid effect of H3K27M is a recent high-resolution imaging study suggesting that although H3K27M does not change the amount of PRC2 bound to chromatin, it can increase the time needed for PRC2 to find and select its target chromatin loci (*31*). An implication of these studies is that PRC2 is not stable on H3K27M chromatin, an effect with striking resemblance to the recently proposed “hit and run” hypothesis of PRC2 where core PRC2 devoid of “stabilizing” components such as MTF2 and JARID2 is less stable on chromatin (*27*). These findings therefore set the stage for future proteomic studies designed to test whether it is indeed decreased association of PRC2 with these stabilizing co-factors or other more direct effects of H3K27M on the PRC2 core complex that drives the transient H3K27M-PRC2 interaction, or alternatively a rapid change in the epigenome by the gain of H3K36me2 (these studies) as well as of H3K27ac (*6*).

One of the most unexpected findings in our study was a second, previously unknown mode of PRC2 inhibition that apparently persists even after its dissociation from H3K27M (Fig. 4). Importantly, this mode of inhibition was more severe as the concentration of SAM increased, suggesting that the PRC2 released from K27M might have a defect in SAM turnover, thereby delaying catalysis (*9*). The other possibility is that H3K27M might inhibit EZH2 automethylation (*32, 33*), eventually influencing the activity of PRC2 on chromatin. Recent studies from our group and others revealed novel automethylation sites on EZH2 and highlighted the impact of EZH2 automethylation on PRC2 activity (*32, 33*). It is intriguing to speculate that EZH2 automethylation could stably change the conformation of the PRC2 complex thereby altering its activity on chromatin. Seemingly in line with a lasting effect of H3K27M on PRC2, is a recent study showing that while the levels of PRC2 bound to chromatin do not change, the search time of PRC2 for chromatin is increased in cells that express H3K27M (*31*). While other variables such as lower H3K27me3 levels may affect PRC2 search time, it is consistent with our findings that there may be a lasting change in PRC2 following its dissociation from H3K27M-containing chromatin that affects its later function.

Our mechanistic studies were also able to give insight into the paradoxical observation that despite a wide-spread loss in H3K27me3, there are a smaller number of loci that actually gain H3K27me3 in DIPG at steady state (*4, 5, 7, 12*). We showed that H3K27M has a strong inhibitory effect on the allosteric activated form of PRC2 (Fig. 5). To put our finding in the context of the current literature, the initial step of allosteric activation of PRC2 is its binding to H3K27me3, the terminal enzymatic product of PRC2, through its essential EED subunit.

The result is a conformational change in EZH2 that stabilizes its catalytic SET domain leading to a substantial enhancement of its HMT activity (*9, 34*). This positive feedback loop facilitates the spreading of H3K27me2/−3 from PRC2 recruitment sites (*27, 35*). Here, we observed that allosterically activated PRC2 preferentially binds to H3K27M nucleosomes (Fig. 5C), an effect similar to one previously reported using structural studies (*9*). Yet, we found that the strongest inhibitory effect of H3K27M was on the allosterically activated PRC2, implying a blockade of the positive feedback activation loop. The net effect is that PRC2 is still recruited to strong polycomb foci in H3K27M cells. However, PRC2 is not allosterically activated and thus, H3K27me3 does not spread, leading to the accumulation of H3K27me3 at strong PRC2 foci. Together, these modes of PRC2 inhibition shed light onto how gains of H3K27me3 can occur at select regions despite the more global loss in PRC2 activity.

In the context of the broader chromatin landscape in H3K27M, we showed both large gains in histone acetylation (fig. S5) as previously reported (*3, 6, 12*), as well as the novel and robust increase in another transcriptionally active histone modification, H3K36me2 (Fig. 6). Based on recent studies suggesting that disrupting the balance of PRC2 activity and H3K36 methylation levels can drive oncogenesis, we speculate that high H3K36me2 might represent a core feature of H3K27M DIPG (*30*). It is worth noting that given the heterogeneity present within K27M DIPG compared to WT glioma, more work is needed to better understand the precise contributions of H3K36me2 to transcriptional programs that might drive oncogenesis specifically in K27M DIPG. Such studies of the downstream effects of H3K36me2 will likely be fruitful since readers of this modification as well as the enzymes that place this modification are potentially drugable therapeutic vulnerabilities in DIPG. Indeed, targeting oncogenic chromatin features, such as residual PRC2 activity as well as the gain of histone acetylation marks and their readers, have been shown to decrease the hallmarks of DIPG growth and aggression in preclinical models (*2, 5, 6, 29, 36*). Therefore, combining biochemistry with mechanistic studies, proteomics, and DIPG disease model systems promises to not only shed light on the underlying biology, but also to reveal novel targets, alone or in combination, for therapeutic intervention in this otherwise intractable disease.

## MATERIALS AND METHODS

### Cell Culture Models and Tissue Preparation

A complete list of cell lines and *in vivo* H3K27M model systems can be found in **Table S1**. Culture methods are described briefly, below.

#### Mouse Stem Cells and Motor Neuron Differentiation

Mouse E14 ESC were cultured as described previously (*17*). Differentiation to cervical motor neurons is detailed in Narendra et al., 2015, but briefly, ESC were plated at low density in the absence of mitogenic factors to form embryoid bodies. Two days later, media was supplemented with 1 μM retinoic acid and 0.5 μM smoothened agonist. Cells were harvested 4 days later with one media replenishment at 2 days.

#### Human Brian Tumor Models

Patient-derived glioma lines, including DIPG lines (see **Table S1** for source), were cultured in tumor stem cell media as described (*37, 38*). Human neural stem cells were derived and cultured as described previously (*39*). NYU patient derived tumor cell line generation has been approved by the NYU Institutional Review Board (IRB# S15-01228). Pathological diagnoses of primary tumors from which cell lines were derived, were performed by a board certified neuropathologist (M.S.) and molecular characteristics were established at an NYU CLIA certified molecular pathology laboratory. Furthermore, relevant tumor sample collection and molecular profiling were approved by the NYU Institutional Review Board (IRB#: S12-00865 and S14-00948, respectively). All human tumor samples were reviewed and diagnosis performed by a board certified neuropathologist (M.S.) in a CLIA certified pathology laboratory.

### Generation of Isogenic Cell Lines

#### mESC Model

The H3K27M mutation was introduced into the H3F3A locus through CRISPR-Cas9 engineering as described in Narendra et al., 2015. Briefly, gRNA directed at H3F3A were designed using the http://crispr.mit.edu/ design tool and cloned into the SpCas9-2A-GFP vector (Addgene Plasmid #48138). mESC were co-transfected with the Cas9, guide RNA and donor DNA template to promote homology directed repair resulting in an M substitution at K27 of H3F3A (i.e., AAG to ATG). Cells containing the Cas9 vector were selected by GFP+ FACS sorting and single cells were plated at low density. Individual colonies were picked 5-7 days later. Individual clones bearing the mutation were harvested and genotyped using Sanger sequencing. Colonies positive for the H3F3A K27M substitution were passaged at least 5 times to confirm purity of the population.

#### 293 TREX Model

Cells were cultured using standard media (DMEM + FBS + metabolic support). Stable cell lines in which the expression of the integrated transgene, either H3.3K27WT or H3.3K27M, under the control of doxycycline were generated by transfecting pINTO-NFH vector (*40*). Stable cell lines were selected using 200 μg/mL zeocin and 10 μg/mL blasticidin selectable markers encoded in the pINTO-NFH vector. Stable integration was subsequently confirmed by Sanger sequencing. To induce expression of the H3.3K27WT or H3.3K27M transgenes, 1 g/mL of doxycycline was added at the appropriate time point prior to harvesting.

A derivative of HEK-293 TREX suitable for PRC2 purification was generated as follows. A 6xHIS-tagged EZH2 cDNA and FLAG-tagged EED cDNA were subcloned into the pLVX-EF1alpha-IRES-mCherry lentiviral vector (Clontech). Cells stably expressing both 6xHIS-EZH2 and FLAG-EED cDNAs were generated by lentiviral transduction. For the production of viral particles, lentiviral vectors were co-transfected with pcREV, BH10, and pVSV-G into HEK-293 TREX cells. The virus-containing medium was collected 48 hr after transfection. The viruses carrying 6xHIS-EZH2 or Flag-EED were mixed in a 1:1 ratio prior to infection of 293 TREX cells. Polybrene was added to the viral medium at a concentration of 8 ug/mL. Infected cells were selected using mCherry expression by FACS sorting.

### Purification of recombinant proteins

To purify the human PRC2 core complex, FLAG-tagged-EED, 6xHIS-tagged-EZH2, SUZ12, and RBAP48 were cloned into the baculovirus expression vector pFASTBac1 (Invitrogen). Recombinant PRC2 was produced in SF9 cells grown in SF-900 III SFM (Invitrogen). After 60 hr of infection, SF9 cells were resuspended in BC150 (25 mM Hepes-NaOH, pH 7.8, 1 mM EDTA, 150 mM NaCl, 10% glycerol and 1 mM DTT) with 0.1% NP40 and protease inhibitors (1 mM PMSF, 1 mM Benzamidine, 1 μg/ml Pepstatin A, 1 μg/ml Leupeptin and 1 μg/ml Aprotinin). Cells were lysed by sonication (Fisher Sonic Dismembrator model 100), and PRC2 complexes were purified through Ni-NTA agarose beads (Qiagen), FLAG-M2 agarose beads (Sigma), and Q Sepharose beads (GE healthcare).

### Purification of PRC2 from 293 TREX

After expressing H3.3K27M or H3.3K27WT using 1 μg/mL of doxycycline for 72 hr, cells grown in 40×15 cm plates were harvested for each condition. Nuclei were purified using a standard hypotonic lysis approach and nuclear extract was prepared using BC400 buffer (20 mM Tris, pH 7.9, 400 mM KCl, 0.2 mM EDTA, 0.5 mM DTT, 20% glycerol, 0.05% NP-40, 1 mM PMSF, 1 mM Benzamidine, 1 μg/ml Pepstatin A, 1 μg/ml Leupeptin and 1 μg/ml Aprotinin). The first purification step utilized Ni-NTA-agarose beads to purify EZH2 which had a 6xHis tag, followed by Q-Sepharose. The eluate was then further purified using FLAG-M2 agarose beads to isolate the complex containing FLAG-tagged EED, followed by another Q-Sepharose. PRC2 was eluted in BC500 (20 mM Hepes NaOH, pH 7.8, 500 mM NaCl, 0.02% NP-40, 10% glycerol, 1 mM PMSF and 1 mM Benzamidine). Components of purified PRC2 were visualized by silver staining after SDS-PAGE and its activity gauged in HMT assays. PRC2 was also purified from 293 cells containing FLAG-tagged EED, but not 6xHis tagged EZH2 using FLAG-M2 agarose and Q-Sepharose.

### Nucleosome reconstitution

Purification of recombinant histones, refolding recombinant octamers, and reconstitution of nucleosomes were generated as described previously (*25*).

### Histone methyltransferase assay (HMT)

Standard HMT assays were performed as described (*25*) unless otherwise stated in the figure legends. Briefly, the reaction was performed in a total volume of 15 μL of HMT buffer (50 mM Tris-HCl, pH 8.5, 5 mM MgCl2, and 4 mM DTT) with the indicated amounts of recombinant human PRC2 complex (EZH2, EED, SUZ12, and RBAP48), 500 nM of 3H-labeled S-Adenosylmethionine (Perkin Elmer). Specific conditions related to the main Figures are presented below with the Supplemental figure detailed information provided.

All reaction mixtures were incubated for 60 min at 30 °C and stopped by the addition of 4 μL SDS buffer (0.2 M Tris-HCl, pH 6.8, 20% glycerol, 10% SDS, 10 mM β-mercaptoethanol and 0.05% Bromophenol blue). After HMT reactions, samples were incubated for 5 min at 95 °C and resolved on SDS-PAGE gels. The gels were then subjected to Coomassie blue staining for protein visualization or wet transfer of proteins to 0.45 μm PVDF membranes (Millipore). The radioactive signals were detected by exposure on autoradiography films (Denville scientific).

### PRC2 Quantitation by Mass-spectrometry

#### Cell Lysate Preparation

Cells were harvested by counting each cell line in quadruplicate and resuspended in a known volume of lysis buffer (8 M Urea, 1 % CHAPS and 20 mM HEPES, pH 8.0). The suspension was pulsed 6x with a sonic dismembrator on ice and rotated end over end for 1 hr at 4 oC. DNA was pelleted and cell lysate was prepared for mass-spec analysis.

Cell lysates were prepared using the filter-aided sample preparation (FASP) method. Briefly, 50 - 200 μg of protein per sample, at 1 – 10 μg/μl concentration, was reduced with DTT (final concentration 3 mM) at 57 oC for 1 hr and loaded onto a Microcon 30 kDa centrifugal filter unit pre-equilibrated with 200 μl of FASP buffer A (8 M urea and 0.1 M Tris HCl, pH 7.8) (Millipore). Following three washes with FASP buffer A, lysates were alkylated on filter with 50 mM iodoacetamide, 45 min in the dark. Filter bound lysates were then washed 3 times with FASP buffer A, 3 times with FASP buffer B (100 mM ammonium bicarbonate, pH 7.8) and digested overnight at RT with trypsin (Promega) at a 1:100 ratio of enzyme to protein. Peptides were eluted twice with 0.5 M NaCl. Tryptic peptides were desalted using an Ultra-Micro Spin Column, C18, (Harvard Apparatus) and the desalted peptide mixture was concentrated in a SpeedVac concentrator.

#### Targeted Mass Spectrometry Assay Development

Peptides for use in this assay were selected based on a series of experiments. First, we compiled data from the Global Proteome Machine Database (GPMDB) for each of the three proteins (*41*). GPMDB is a repository of the results of mass spectrometry based proteomics experiments. The data collected included the peptides identified, the charge states identified, their uniqueness to the protein, and the number of times each was identified. The most often identified unique peptides and charge states of EED, EZH2, and SUZ12 were compared to the results of the recombinant analysis to generate a list of approximately 5 candidate peptides per protein.

Next, an equal molar trypsin digested mixture of the three recombinant mouse PRC2 proteins (EED, EZH2, and SUZ12) was analyzed on the Q Exactive mass spectrometer (Thermo Scientific) using a data dependent method. A list of identified proteotypic peptides was compiled. This list of peptides was used to generate a target list in the Skyline software, which was used to build the targeted MS2 method. An aliquot of a mESC lysate was analyzed using this method. Raw files were uploaded to the Skyline software. Candidate peptides were evaluated based on the chromatographic peak shape, ionization efficiency, coverage of fragmentation, level of interference from other peptides within the isolation window and the presence of product ions that differentiate the endogenous and heavy labeled peptide. 2 – 3 of the strongest candidate peptides were custom synthesized (New England Peptide) with an isotopically heavy C-terminal residue.

#### Targeted Mass Spectrometry Analysis

The peptides were spiked into an aliquot of the cell lysate at a level ranging from 100 amoles to 5 fmoles. The peptide mixture was loaded onto an Acclaim PepMap trap column in line with an EASY-Spray 50 cm x 75 μm ID PepMap C18 analytical HPLC column with 2 μm bead size using the auto sampler of an EASY-nLC 1000 HPLC (Thermo Scientific). The peptides were gradient eluted into a Q Exactive (Thermo Scientific) mass spectrometer using a 60 min gradient from 5% to 23% solvent B (Solvent A: 2% acetonitrile, 0.5% acetic acid; Solvent B: 90% acetonitrile, 0.5% acetic acid), followed by 20 min from 23% to 45% solvent B. Solvent B was taken to 100% in 10 min and held at 100% for 20 min. High resolution full MS spectra were acquired with a resolution of 70,000, an AGC target of 1e6, with a maximum ion time of 120 ms and scan range of 300 to 1500 m/z. Following each full MS scan, targeted high resolution HCD MS/MS spectra were acquired based on the inclusion list containing the m/z values of the unlabeled endogenous and stable isotope labeled standard PRC2 peptides. All MS/MS spectra were collected using the following instrument parameters: resolution of 17,500, AGC target of 2e5, maximum ion time of 120 ms, one microscan, 2.0 m/z isolation window, fixed first mass of 150 m/z, and Normalized Collision Energy (NCE) of 27. Peptides not behaving linearly in lysate were excluded from future quantitation.

Quantitative experiments were conducted in two ways: 1) 5 point calibration curve (500 amole to 5 fmole) of standard peptides spiked into a constant amount of lysate or 2) a single point calibration of standard peptides spiked into a constant amount of lysate in triplicate in a concentration close or equal to the endogenous level.

#### Data Analysis and validation

Raw files were uploaded to Skyline (64 bit) software version 3.6.0.10162 for analysis. The top three or four product ions were used to quantify the light (endogenous) and stable isotope-labeled standard peptides. Total product ion area values were calculated for the light and heavy peptides.

When a calibration curve was generated, the total product ion area values for each of the stable isotope-labeled standard peptides was plotted, a linear trendline was fit to the data and the line equation was used to calculate the fmoles of endogenous peptide in the lysate. The average, standard deviation and the coefficient of variance across the triplicates is reported for all lysates.

To calculate the number of each PRC2 subunit (EZH2, EED and SUZ12) per cell, the average fmole for each peptide quantified of that subunit was computed and converted to number of molecules using Avogadro’s number and divided by the number of cells analyzed to obtain molecules per cell. Total PRC2 per cell was calculated by taking the average of all subunits.

A few attempts have been made to infer the number of PRC2 molecules per nucleus using a variety of imaging and enzyme kinetic techniques. Reports vary from numbers as low as 6,000 [(ALL/myeloma;(*42*)] to ~40,000 [(*Drosophila* neuroblast; (*43*)] and as high as 20,006,965 in *Drosophila* salivary gland (*44*). While our methods provided slight variations on these numbers, they are well within the range and ordinal differences reported, adding validity to these various methods of PRC2 quantitation.

### Histone Extraction, Mass-Spectrometry and Analysis of Histone Modifications

Histone extraction, mass-spectrometry and analysis of histone modifications were based on previously published methods (*45, 46*).

### ChIP-seq Experiments

ChIP was performed as described previously (*40*). Briefly, H3.3K27M was induced by 1 ug/mL doxycycline for 72 hr, cross-linked with 1% formaldehyde and harvested following glycine treatment to quench the formaldehyde. Nuclei were extracted, sonicated using a Diagenode Bioruptor to produce chromatin fragments at an average size of 300-500 bp. Chromatin was then precleared with BSA blocked magnetic Dynabeads. Each ChIP was performed with an internal spike-in standard corresponding to 1:200 of cross-linked, sonicated and pre-cleared *Drosophila* chromatin. The *Drosophila*-specific H2A.V antibody was also added at ~1:20 of the primary ChIP antibody. The immunoprecipitation was performed using antibodies to the appropriate protein (**Table S2**) and 30 uL of BSA-blocked Dynabeads. Immunoprecipitated chromatin bound to the beads was then washed in RIPA buffer. Chromatin was de-crosslinked and proteins removed by proteinase K. DNA libraries were constructed, barcoded as in (*17*) and subjected to SR50 sequencing on an Illumina HiSeq2500 or Illumina NextSeq 500.

### ChIP-seq Data Processing

Quality of sequencing was assessed with FastQC v0.11.4 (http://www.bioinformatics.babraham.ac.uk/projects/fastqc/). Reads having less than 80% of quality scores above 25 were removed with NGSQCToolkit v2.3.3 (*47*) using the command *IlluQC.pl -se $file_path N A -l $percent - s $threshold -p $nb_cpu -o $out_path*. Reads were aligned to the human hg19 reference genome from Illumina igenomes UCSC collection using bowtie (*48*) v1.0.0 allowing 3 mismatches and keeping uniquely aligned reads (*bowtie -q -v 3 -p $nb_cpu -m 1 -k 1 --best --sam --seed 1 $bowtie_index_path “$file_path”*). Sam ouputs were converted to Bam with Samtools (*49*) v1.0.6 (*samtools view -S -b $file_path -o $out_path/$file_name.bam*) and sorted with Picard tools v1.88 (http://broadinstitute.github.io/picard: *java -jar SortSam.jar SO=coordinate I=$input_bam O=$output_bam*). Data were further processed with Pasha (*50*) v0.99.21 with the following parameters: WIGvs = TRUE, incrArtefactThrEvery = 7000000, elongationSize = NA. Fixed steps wiggle files were converted to bigwigs with the script wigToBigWig available on the UCSC Genome Browser website *(http://hgdownload.soe.ucsc.edu/admin/exe/)*.

Spike-in normalization was achieved using *Drosophila Melanogaster* DNA aligned to Illumina igenomes dm3 as mentioned above (*51*). Endogenous and exogenous scaling factors were computed from the bam files (1/(number_mapped_reads/1000000)). Endogenous scaling factors were applied to the data before input subtraction (without scaling). The RPM normalization was then reversed before scaling with exogenous factors. The scripts used to perform spike-in scaling have been integrated in a bioconductor package ChIPSeqSpike (https://bioconductor.org/packages/3.7/bioc/html/ChIPSeqSpike.html).

### Domain selection and peak calling

The different domains were identified in 293 TREX cells at all time points with hiddenDomains 3.0 (*52*), with the command ‘*hiddenDomains -g $chromInfo_file -b $bin_size -t $expBam -c $inputbam -o $outputFilePrefix’*. A window size of 50 or 100 kb was used (**Fig. 2 3 and S3** for example). To more precisely identify overlap between ChIPs within these domains, peaks were detected within the domain as noted in the text and figure legends. Briefly, peaks were detected using SeqMonk v1.41 (http://www.bioinformatics.babraham.ac.uk/projects/seqmonk/) merging biological replicates using a 1e-05 cutoff and were then filtered to identify only peaks contained within the domains and transformed to the heatmap representation noted below (**Fig. 2&3**).

To retrieve the common K27me3 peaks by overlap in DIPG cells (**Fig. 5**), MACS2 v2.1.0 was used (*macs2 callpeak -t $bam_file_vector -c $input_file_vector -n $experiment_name --outdir $output_folder_nomodel_broad -f $format -g $genome_size -s $tag_size --nomodel --extsize $elongation_size --keep-dup $artefact_threshold --broad --broad-cutoff 1e-05*).

### Metagene profiles and heatmaps

Metagene profiles and heatmap were largely generated with the bioconductor package seqplots (*53*) v1.8.1 in command line. Mean, 95% confidence intervals and standard error are indicated on the profiles in the figures.

Heatmaps for **Fig. 2 3** and as well as quantitative trend plots for **Fig. 5**(1,727 or all refseq genes) were generated with Deeptools v2.3.3 (*54*)

#### Data availability

The data will be submitted to GEO prior to publication.

### Western Blot

Western blots were performed largely as in (*25, 27*) with Bis-Tris resolving gels and transfer to 0.45 μm PVDF membranes. The antibodies used are indicated in the text and **Table S2**.

### General Quantitation and Data Analysis

Band intensity for western blots and autoradiography experiments were quantitated after normalization using Image J. Where appropriate, simple *a priori* hypotheses were tested by ANOVA followed by Fisher’s LSD. Experimental alpha was set equal to 0.05 for all experiments.

## Acknowledgments

**General**

We thank Drs. L. Vales and K.-J. Armache for their critical guidance throughout the studies and their critical reading of the manuscript. We thank Drs. P. Lee, S. Krishnan, E. Campos, A. Rojas, V. Narendra as well as other past and current Reinberg lab members for their discussion as the work was in progress. We are also grateful to D. Hernandez and N. Jahan for their technical assistance and to Dr. Michelle Monje and Dr. Esther Hulleman for kindly providing DIPG/glioma cell lines that were integral to this work.

## Funding

the NYU Flow Cytometry Core, NYU Proteomics Laboratory and NYU Genome Technology Center are partially supported by the NYU School of Medicine and the Laura and Isaac Perlmutter Cancer Center support grant, NCI (P30CA016087). The work was supported by grants to D.R. from NIH (R01CA199652), HHMI and the Making Headway Foundation (189290). A grant from the Hyundai Hope on Wheels Foundation to M.S. and D.R. partially supported this work. Molecular profiling of human samples was in part supported by grants from the Friedberg Charitable Foundation, the Rachel Molly Markoff Foundation and the Sohn Conference Foundation to M.S. J.M.S was a Simons Foundation’s Junior Fellow and was also supported by an NIH (K99AA024837) grant. J-R.Y. is supported by the American Cancer Society (PF-17-035-01). P.V. was supported by the Empire State Training Program in Stem Cell Research (NYSTEM, contract no. C026880).

## Author contributions

J.M.S., C-H.L., P.V. and D.R. conceptualized and designed the study. J.M.S., C.-H.L., P.V., J.-R.Y., G.L., O.O., and F.S. conducted the experiments. N.D. and R.S.M. performed bioinformatics analyses. J.R.C. and B.U. performed mass spectrometry analyses. A.S.M., N.S.B., D.G.P., M.A.K., M.S. provided guidance on brain tumor models as well as provided cell lines/banked tumors. J.M.S., C.-H.L., R.S.M., N.D. and D.R wrote the manuscript.

## Competing interests

Authors declare no competing interests.

## Data and materials availability

Data and relevant computations for the proteomics work is presented in the Supplementary Data files. Genome sequencing results is uploaded to GEO (GSE118954). All other data and information not available in the main text or supplement is available upon request.

## SUPPLEMENTARY MATERIALS

**Table S1.** Tumor samples and cell lines used in this study.

| Cell Line/Tumor | Origin/Tumor Type | Histone Status | Age | Manipulation | Prior Characterization |
| --- | --- | --- | --- | --- | --- |
| DIPG-VI | DIPG | H3F3A K27M | 7 | N/A | Caretti et al., 2014 |
| DIPG-XIII | DIPG | H3F3A K27M | 6 | N/A | Caretti et al., 2014 |
| DIPG-N | DIPG | HIST1H3C K27M | 5 | N/A | This study |
| DIPG-IV | DIPG | HIST1H3B K27M | 2 | N/A | Caretti et al., 2014 |
| DIPG VUMC (DIPG X) | DIPG | WT | 12 | N/A | Nagaraja et al., 2017 |
| DIPG-Frozen | DIPG | WT | 5 | N/A | This study |
| pcGBM2 | Glioblastoma-Cortex | WT | 15 | N/A | Venkatesh et al., 2015 |
| SF188 | Glioblastoma-Cortex | WT | 8 | N/A | Bax et al., 2009 |
| GBML20 | Glioblastoma-Cortex | WT | 89 | N/A | Bayin et al., 2017 |
| GBML33 | Glioblastoma-Cortex | WT | 45 | N/A | Bayin et al., 2017 |
| MPNST (sNF96.2) | MPNST (schwannoma) | WT | 28 | N/A | DeRaet et al., 2014 |
| hAstrocytes | Normal | WT | N/A | N/A | LONZA |
| hNeural Stem Cells | Normal | WT | N/A | N/A | Modrek et al., 2017 |
| 293 TREX H3K27M | 293 TREX | H3.3K27M | N/A | pINTO-NFH H3.3K27M | This study |
| 293 TREX H3K27M<br>w/PRC2 tag | 293 TREX | H3.3K27M | N/A | pINTO-NFH H3.3K27M<br>pLVX-EF1alpha-IRES-mCherry EED-FLAG<br>pLVX-EF1alpha-IRES-mCherry EZH2-6XHis | This study |
| 293 TREX H3K27WT | 293 TREX | H3.3K27WT | N/A | pINTO-NFH H3.3K27WT | This study |
| 293 TREX H3K27WT<br>w/PRC2 Tag | 293 TREX | H3.3K27WT | N/A | pINTO-NFH H3.3K27WT<br>pLVX-EF1alpha-IRES-mCherry EED-FLAG<br>pLVX-EF1alpha-IRES-mCherry EZH2-6XHis | This study |
| mESC H3F3AK27M | E14 mEmbryonic Stem Cell | H3F3A K27M | N/A | CRISPR Cas9 at H3F3A Locus | This study |
| mESC H3F3AK27WT | E14 mEmbryonic Stem Cell | WT | N/A | CRISPR Cas9 Control | This study |

**Table S2.** Antibodies and relevant ChIP conditions used in this study

| Antibodies | SOURCE | IDENTIFIER | ChIP Experimental Notes |
| --- | --- | --- | --- |
| EED | Reinberg Lab | N/A |  |
| EZH2 (D2C9) | Cell Signaling Technology | Cat# 5246,<br>RRID:AB_10694683 | ChIP: 150µg Chromatin<br>per 8µL antibody |
| SUZ12 (D39F6) | Cell Signaling Technology | Cat# 3737BF,<br>RRID:AB_2616021 |  |
| H3K27me2 (D18C8) | Cell Signaling Technology | Cat# 9728 |  |
| H3K27me3 (C36B11) | Cell Signaling Technology | Cat# 9733<br>RRID:AB_2616019 | ChIP: 50µg Chromatin<br>per 4µL antibody |
| H3K36me2 (C75H12) | Cell Signaling Technology | Cat# 2901 | ChIP: 50µg Chromatin<br>per 4µL antibody |
| HA (16B12) | Biolegend | Cat# 901502 |  |
| HA (ChIP-grade) | Abcam | Cat# ab9110 | ChIP: 50µg Chromatin<br>per 4µL antibody |
| FLAG | Sigma-Aldrich | Cat# F1804,<br>RRID:AB_262044 |  |
| H3 | Abcam | Cat# ab1791,<br>RRID:AB_302613 |  |
| Islet 1/2 | UNIV. OF IOWA/Developmental<br>Studies Hybridoma bank | 39_4D5 |  |
| GAPDH | GeneTex | Cat# GTX627408,<br>RRID:AB_11174761 |  |

**Supplemental Figure 1.**
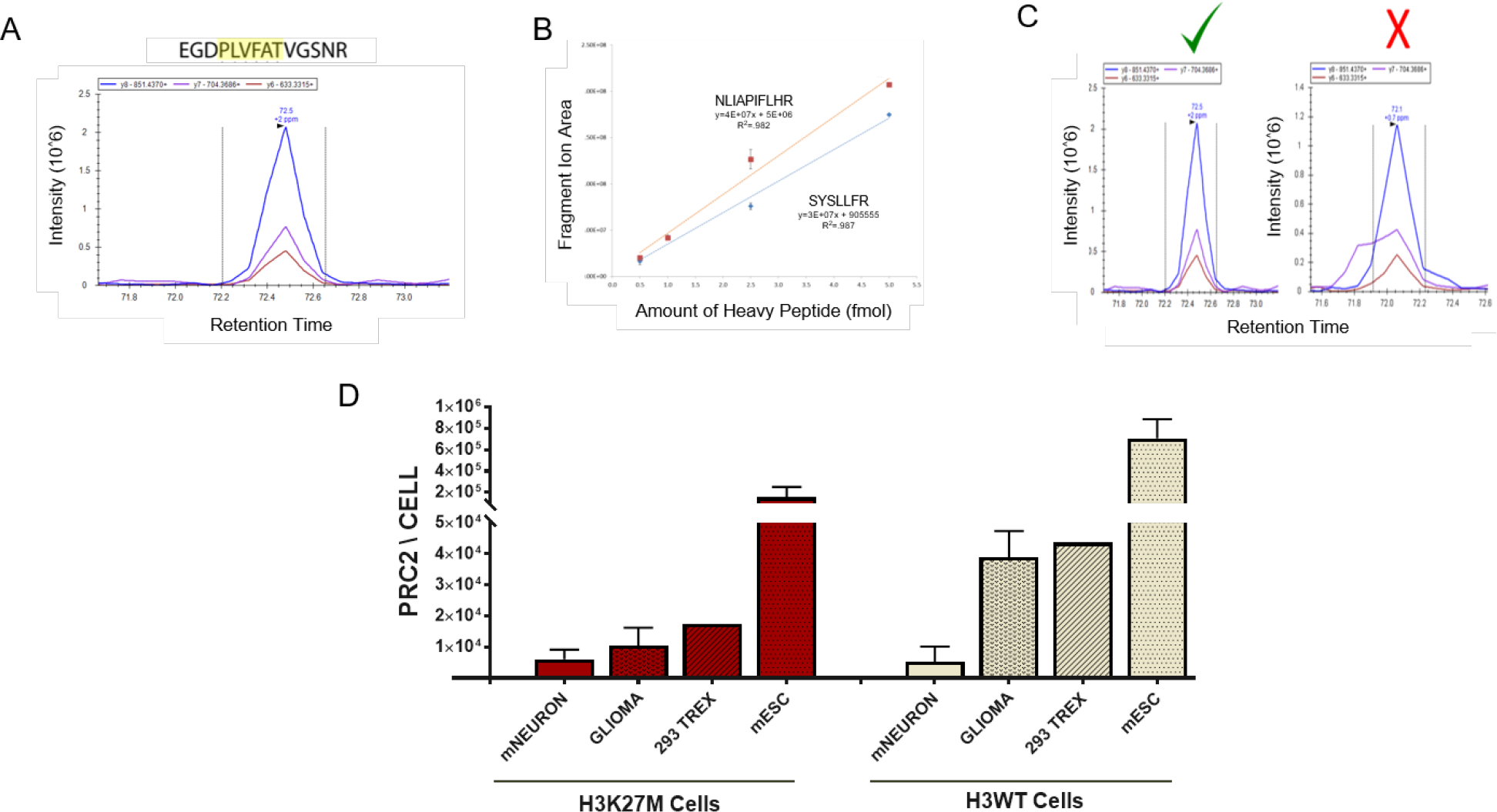
PRC2 Quantitation Across Cell Types. **(A)** Frequently detected, proteotypic peptides for each PRC2 component were selected from the Global Proteome Machine Database and cross-referenced to tryptic peptides observed in recombinant PRC2. **(B)** Heavy labeled peptides were spiked into cell lysate at different amounts to ensure linear behavior. Representative quality control of an example of two validated SUZ12 peptides are shown. **(C)** Heavy labeled peptides were selected for minimal interference from the extract matrix. Peptides passing these quality control steps were used in final quantitation (representative SUZ12 peptide is shown as an example). **(D)** PRC2 quantitation in all cells. In the isogenic cell lines, sample size were; H3K27M mNEURON *n=2*, H3K27M 293 TREX *n=1*, H3K27M mESC *n=3*, H3WT mNEURON *n=2*, H3WT 293 TREX *n=1*, H3WT mESC *n=4*. In the glioma cell lines the sample sizes and specific cell lines used were as follows; H3K27M Glioma (DIPG XIII, DIPG N, DIPG IV) and H3WT Glioma *n=3* (pcGBM2, GBML20, DIPG X). Data Plotted as Mean±SEM.

**Supplemental Figure 2.**
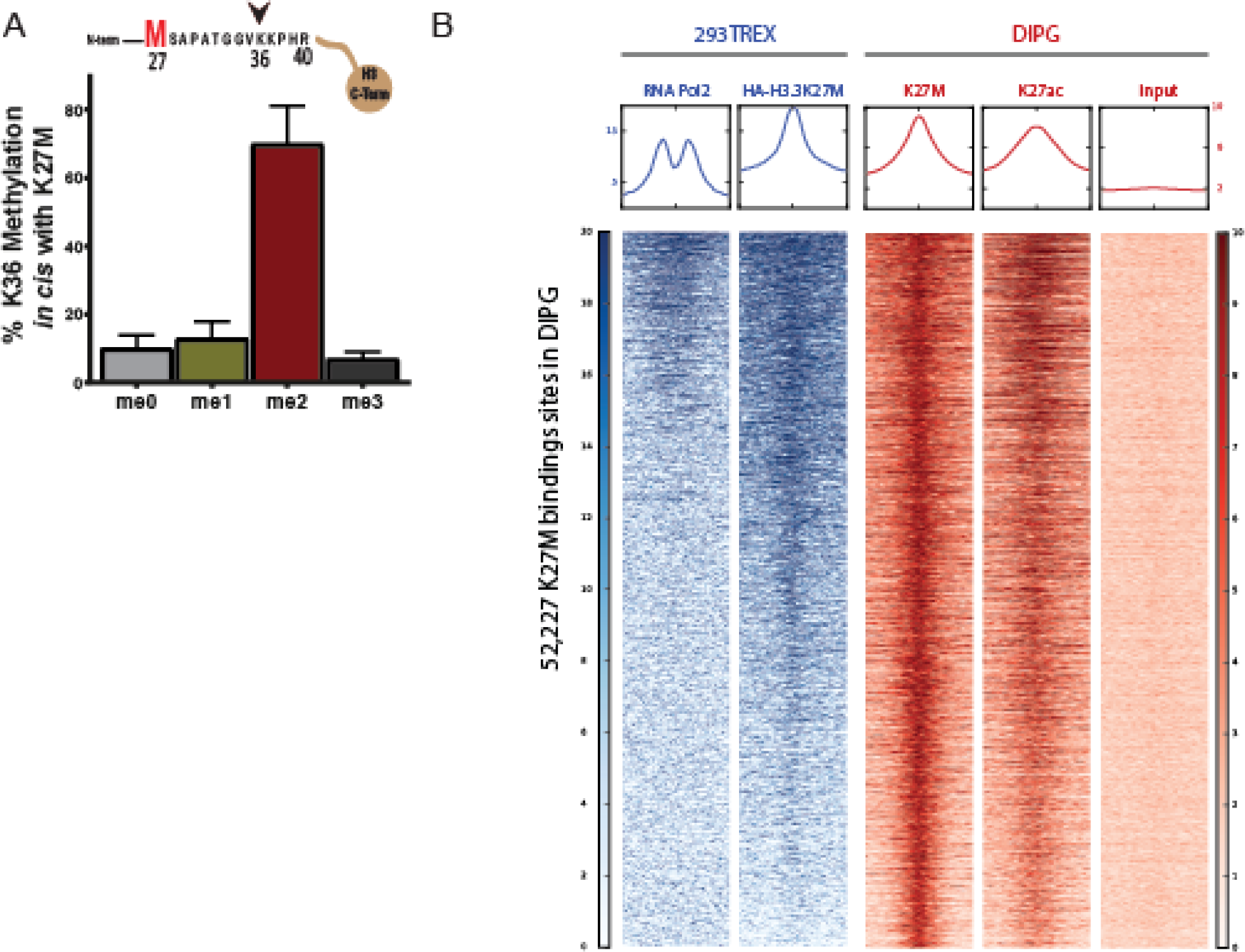
H3K36me2 is the predominant modification adjacent to K27M and K27M is deposited primarily into active regions. **(A)** Quantitative histone MS graphs indicating that K36me2 is the predominant methylation adjacent to K27M (in cis; *n*=4 H3K27M DIPG). Data Plotted as Mean±SEM. **(B)** Heatmap representation of 293 TREX (blue) and DIPG [red - GSE78801 (*6*)] ChIP-seq data for Pol2 (*40*), HA (H3.3K27M after 6 hr of induction; this study), K27M, K27ac and Input (*6*). In all cases, read density is displayed 3 kb from each side of the center of the binding site sorted by enrichment of HA.

**Supplemental Figure 3.**
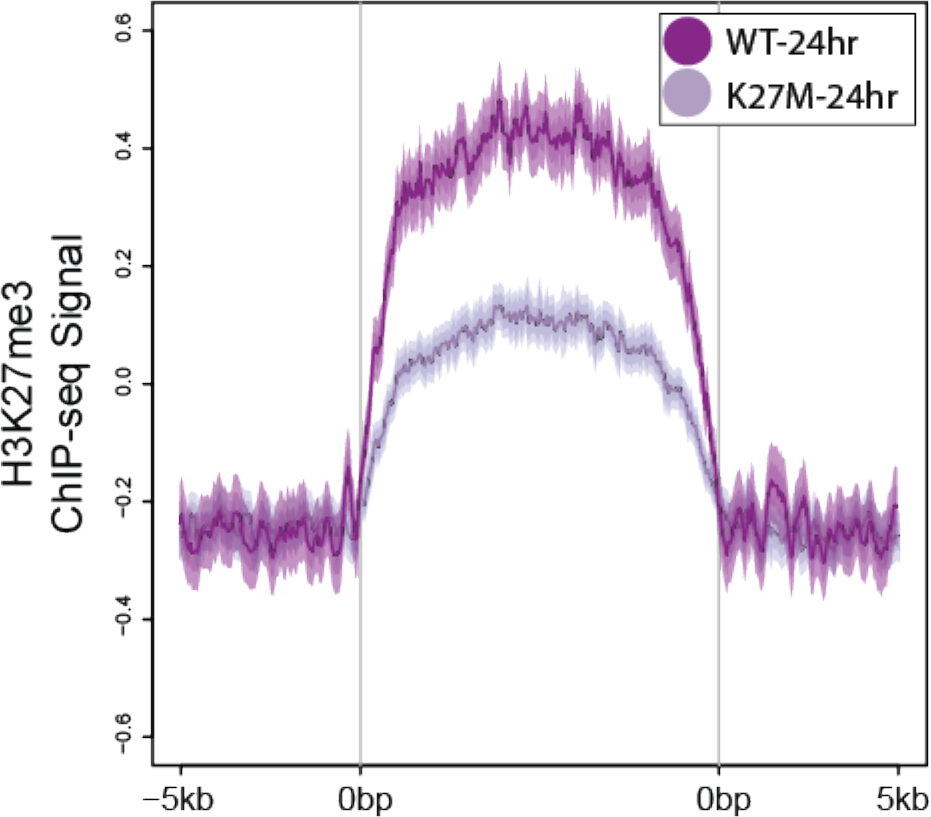
The level of H3K27me3 at PRC2 common targeting domain is reduced in H3K27M cells. 616 large domains that showed PRC2 (EZH2) enrichment across all timepoints were identified and noted as “Common Domains.” For visualization, H3K27me3 is plotted on the domain for the 24 hr timepoint after applying a sliding window to depict its relative size.

**Supplemental Figure 4.**
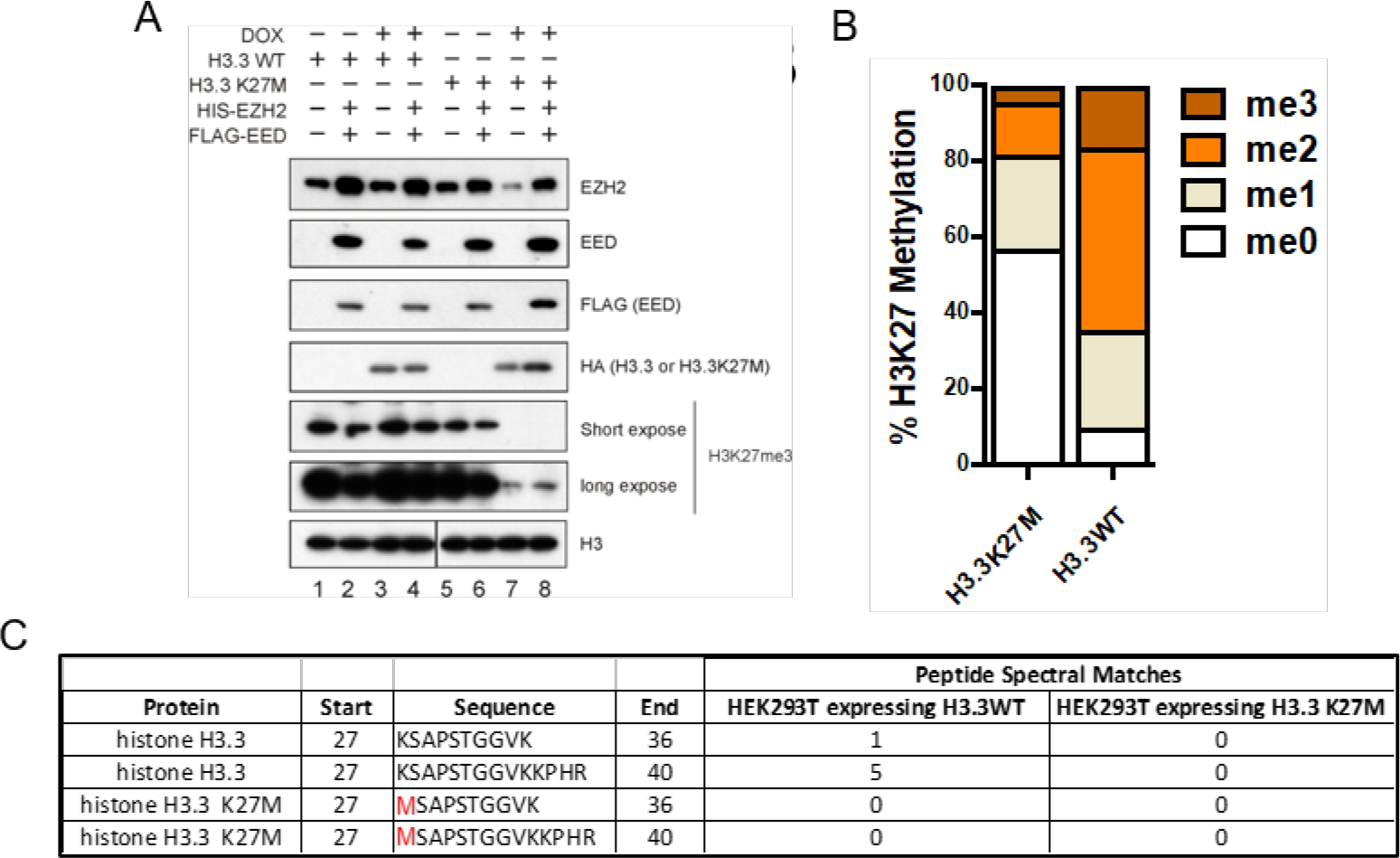
Cell line generation and characterization of PRC2 purified from H3.3K27M or H3.3K27WT cells. **(A)** Validation of cell lines generated for results in **Figure 4**. 293 TREX cells with doxycycline inducible H3.3K27M- or H3.3K27WT-bearing expression vectors and encoding 6xHIS-Tag on EZH2 and FLAG-tag on EED were probed for histones and PRC2 subunits. Expression of FLAG-tagged EED and HIS-tagged EZH2 was confirmed by western blot on cell lysates, as described. Induction was performed for 3 days with 1 μg/mL doxycycline showing the expression of the H3.3WT or H3.3K27M and a robust decrease in H3K27me3 in 293 TREX cells expressing H3.3K27M. **(B)** Mass-spectrometry confirmed the robust drop in H3K27me2-3 levels following 3 days of expressing H3.3K27M vs. H3K27WT. **(C)** Mass-spectrometry indicating that H3K27M histone peptide was not detectable after purification of PRC2 (see *Fig. 4A*).

**Supplemental Figure 5.**
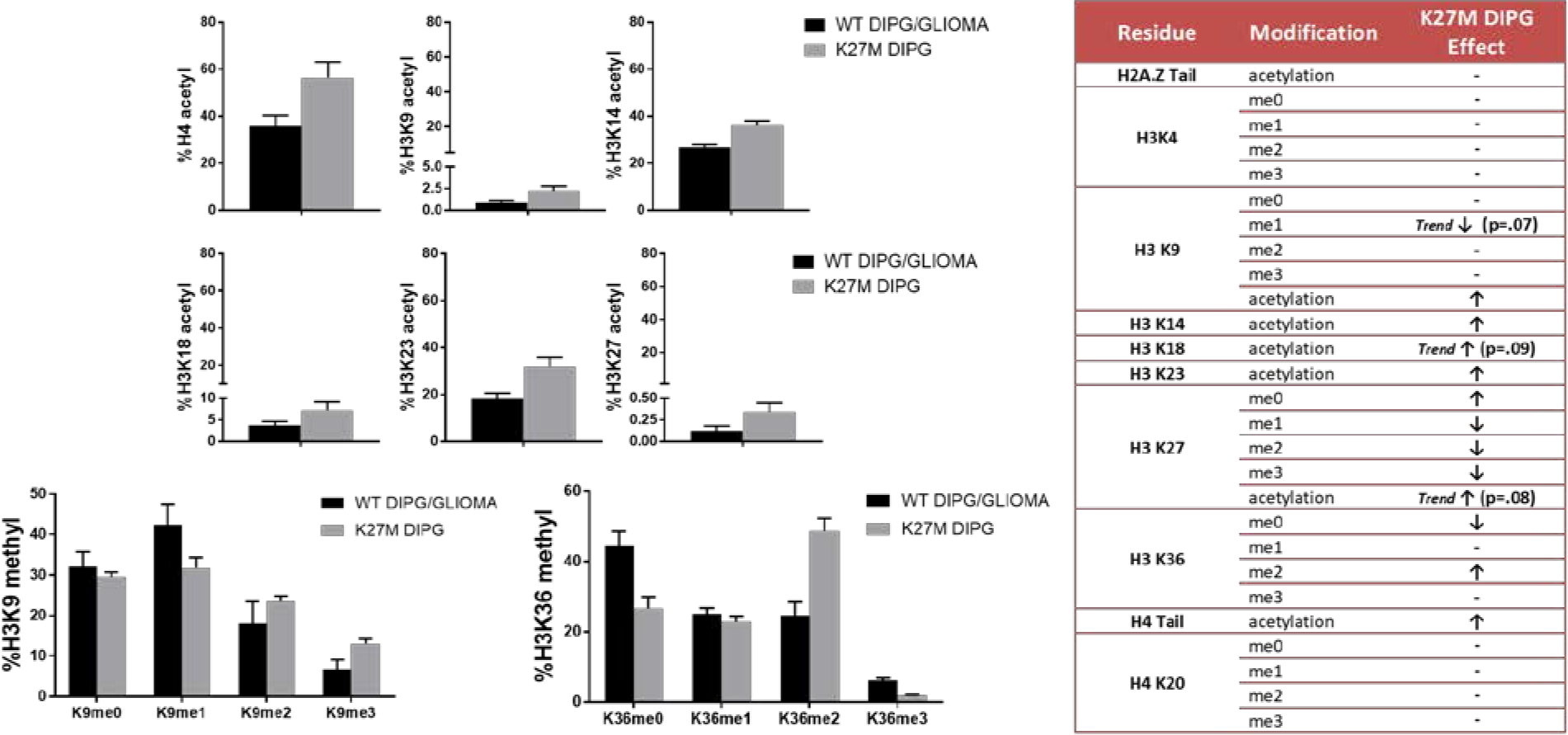
Alterations in Histone Modifications in H3K27M DIPG. Quantitative histone modification mass-spectrometry for modifications significantly altered in K27M-bearing DIPGs (detailed in *Materials and Methods*). **WT DIPG/GLIOMA** (H3K27WT cortical glioma *n=4*, H3K27WT DIPG *n=2)*, **K27M DIPG** (*n=2* for H3.1K27M and *n=2* for H3.3K27M). Data Plotted as Mean±SEM. Summary of significant histone modifications altered in K27M DIPGs is shown in the table on the right (WT samples vs. H3K27M DIPG; Following significant ANOVA for each quantified peptide, Fishers LSD *p <.05* indicates significance, trends noted with *p <.10*). ↑ = increased, ↓ = decreased, -= no change.

**Supplemental Figure 6.**
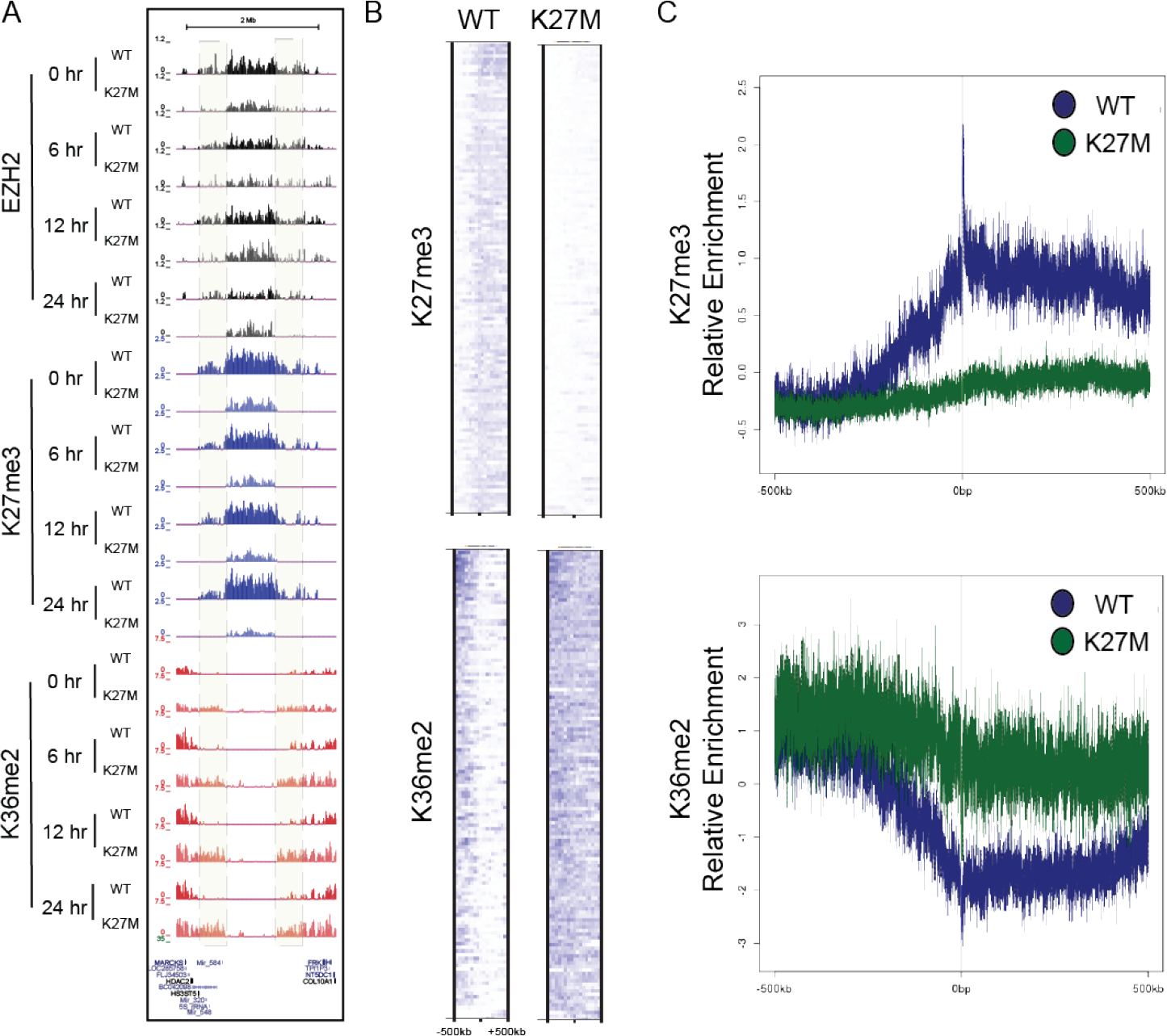
Spreading of K36me2 in the K27M 293 TREX cells. **(A)** ChIP-seq tracks for H3K27me3, H3K36me2 and EZH2 from 293 TREX cells expressing H3.3K27WT (WT) or H3.3K27M (K27M) for 0, 6, 12, or 24 hr. Highlighted are those regions showing increased H3K36me2 over time in H3K27M cells concomitant with a decrease in H3K27me3. **(B)** H3K27me3 domains that are lost in the H3K27M cells across all time points were selected. Heatmaps for H3K27me3 and H3K36me2 were generated by centering and rank ordering the enrichment on those regions. Importantly, only 90 of the 132 sites are represented here to avoid overlap in lines. In doing so, we carefully selected specific cases that represent the most robust domains that lose H3K27me3 in the H3K27M case. While the loss of H3K27me3 occurred in both the 3’ and 5’ direction, we plotted all of those event in the same direction for illustrative purposes. Shown are the corresponding heatmaps for the 24 hr time point. Regions showing loss of H3K27me3 in the H3K27M cells exhibited a gain of H3K36me2. **(C)** Average density profiles corresponding to the heatmap selections indicate that regions which lose H3K27me3 (*top*), show a reciprocal retention of H3K36me2, which likely spreads from the adjacent H3K36me2 domain (*bottom*).

**Supplemental Figure 7.**
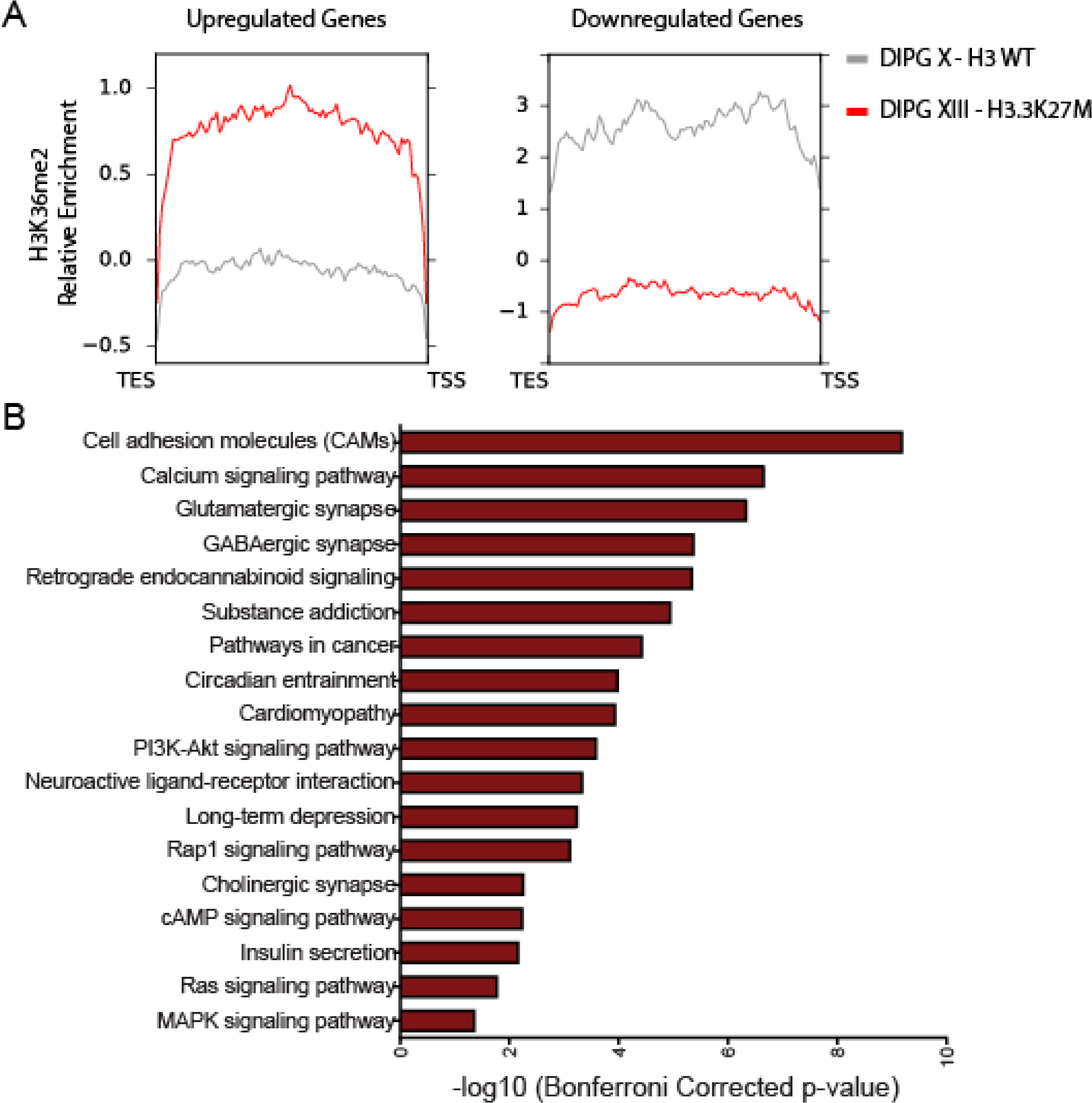
Select transcriptional changes associated with K36me2 in comparable K27M DIPG cell lines. **(A)** RNA-seq was performed on DIPG X (WT) and DIPG XIII (H3.3K27M) and differentially expressed genes were identified using DESeq2. An overall gain of H3K36me2 can be seen on genes that are upregulated (5,297 genes) in the H3.3K27M DIPG XIII cell line. An opposing relationship can be seen for downregulated genes (1,422 genes). **(B)** Ranked KEGG pathway analysis of the upregulated genes (cutoff for inclusion was bonferroni corrected p < .05).

